# Chromosome Conformation Paints Reveal the Role of Lamina Association in Genome Organization and Regulation

**DOI:** 10.1101/122226

**Authors:** Teresa R Luperchio, Michael EG Sauria, Xianrong Wong, Marie-Cécile Gaillard, Peter Tsang, Katja Pekrun, Robert A Ach, N Alice Yamada, James Taylor, Karen L Reddy

**Author notes:** These authors contributed equally to this work (Teresa R Luperchio, Michael EG Sauria). Correspondence should be addressed to JT and KLR.

## Abstract

Non-random, dynamic three-dimensional organization of the nucleus is important for regulation of gene expression. Numerous studies using chromosome conformation capture strategies have uncovered ensemble organizational principles of individual chromosomes, including organization into active (A) and inactive (B) compartments. In addition, large inactive regions of the genome appear to be associated with the nuclear lamina, the so-called Lamina Associated Domains (LADs). However, the interrelationship between overall chromosome conformation and association of domains with the nuclear lamina remains unclear. In particular, the 3D organization of LADs within the context of the entire chromosome has not been investigated. In this study, we describe “chromosome conformation paints” to determine the relationship *in situ* between LAD and non-LAD regions of the genome in single cells. We find that LADs organize into constrained and compact regions at the nuclear lamina, and these findings are supported by an integrated analysis of both DamID and Hi-C data. Using a refined algorithm to identify active (A) and inactive (B) compartments from Hi-C data, we demonstrate that the LADs correspond to the B compartment. We demonstrate that *in situ* single cell chromosome organization is strikingly predicted by integrating both Hi-C and DamID data into a chromosome conformation model. In addition, using the chromosome conformation paints, we demonstrate that LAD (and B-compartment) organization is dependent upon both chromatin state and Lamin A/C. Finally, we demonstrate that small regions within LADs escape the repressive regime at the peripheral zone to interact with the A-compartment and are enriched for both transcription start sites (TSSs) and active enhancers.

## Introduction

DNA is highly organized within the eukaryotic cell nucleus. From sequestration of proteins involved in transcription, RNA processing and DNA repair, into nuclear sub-domains, to emerging evidence of higher order organization of chromatin itself, it has become clear that the nucleus is highly and dynamically organized. Nuclear organization is manifest in the arrangement of chromatin into active and inactive domains (*e.g*. euchromatin and heterochromatin), chromosome territories and sub-chromosomal organizational domains, such as lamina associated domains (LADs) and topologically associated domains (TADs), among others (Albiez et al., 2006; Alexandrova et al., 2003; Boyle et al., 2001; Comings, 1980; Cremer and Cremer, 2001; Dixon et al., 2012; Guelen et al., 2008; Nora et al., 2012). Some architectural domains, such as heterochromatin, preferentially associate with specific spatially distinct regions of the nucleus including the nucleolus and the nuclear periphery. A light microscope and a DNA counterstain is sufficient to observe some of this organization (Boveri, 1914; Heitz, 1929), particularly the partitioning of heterochromatin and euchromatin, but more recent studies have leveraged genome-wide approaches and fluorescence *in situ* hybridization (FISH) to uncover the interrelationship between nuclear organization and genome regulation at different levels of resolution.

Chromosomes appear to occupy distinct regions in the nucleus called chromosome territories (CT), suggesting that each chromosome has a three-dimensional self-interacting organization (Bolzer et al., 2005; Boyle et al., 2001; Cremer and Cremer, 2001; Cremer et al., 2006). These CTs are visible under a microscope by employing whole chromosome-specific DNA probes or “paints” in a FISH assay. Subsequent microscopy-based FISH studies led to the discovery that certain domains within a CT change their relative disposition depending upon activity state, suggesting that intra-chromosomal interactions and organization are tightly linked to gene activity (Cremer and Cremer, 2010). More recently, high-throughput DNA sequencing based approaches, such as Hi-C or similar chromosome conformation capture (3C) based techniques, have been employed to uncover the organization of chromatin into self-interacting domains or modules (Dekker et al., 2002; Dixon et al., 2012; Lieberman-Aiden et al., 2009; Rao et al., 2014). Unlike FISH-based approaches, which target specific loci or chromosomes, these strategies have allowed the genome-wide mapping of all chromatin interactions within the nuclear volume. Two distinct features of these data are the presence of locally self-interacting TADs and a global partitioning into two groups based on longer-range interactions, the so-called chromatin A/B-compartments. The B-compartment largely represents repressed domains while the A-compartment displays robust self-interactions between active regions of the genome(Lieberman-Aiden et al., 2009). The boundaries between these chromatin domains are strongly enriched for architectural proteins, such as CTCF (CCCTC-binding factor) (Dixon et al., 2012; Phillips-Cremins et al., 2013; Rao et al.,2014; Tang et al., 2015). By deconvolving these genome-wide association maps, several attempts have been made to predict a general structure of chromosome domains and folding in the nuclear volume (Hu et al., 2013; Lesne et al., 2014; Segal et al., 2014; Varoquaux et al., 2014; Zhang et al., 2013). Strikingly, such modeling reveals the existence of preferred positions of chromosomal subdomains, including regions enriched for histone H3 lysine 9 di- or trimethylation (H3K9me2/3).

Association of chromatin with the nuclear periphery in particular has been implicated in gene regulation, particularly correlating with gene repression (Finlan et al., 2008; Guelen et al., 2008; Kosak et al., 2002; Luperchio et al., 2014; Peric-Hupkes et al., 2010; Reddy et al., 2008; Zullo et al., 2012). DamID (DNA Adenine Methylation Identification), a genome-wide technique to identify nuclear lamina-proximal chromatin, has allowed identification of Lamina Associated Domains (LADs) (Greil et al., 2006; Vogel et al., 2007). These 100 kilobase (kb) to megabase (Mb) sized domains are enriched for genes that are transcriptionally silent and enriched in histone modifications indicative of facultative heterochromatin, such as H3K9me2/3 and histone H3 lysine 27 trimethylation (H3K27me3) (Guelen et al., 2008; Harr et al., 2015; Wen et al., 2009). Moreover, recent studies have highlighted that both H3K9me2/3 and H3K27me3 are involved in LAD organization (Bian et al., 2013; Harr et al., 2015; Kind et al., 2013; Towbin et al., 2012). LADs are immediately flanked by active promoters, highlighting the stark delineation between repressed LAD domains and adjacent active regions (Guelen et al., 2008). Intriguingly, the architectural protein CTCF, shows enrichment at LAD borders similar to the boundaries between chromatin domains observed by Hi-C (Guelen et al., 2008). In addition, it has been noted that the B-compartment is enriched in LADs and pericentromeric heterochromatin (Lieberman-Aiden et al., 2009; Nora et al., 2012; Schwarzer et al., 2016). However, to date, a detailed exploration of the relationship between LADs and chromosomal sub-domain organization, such as the A/B-compartments, is surprisingly lacking.

How LADs organize at the single cell level remains unknown. One study in a colon cancer derived cell line found that only 30% of regions identified by DamID are lamina-proximal in any single cell (Kind et al., 2013). This would suggest that the organization detected by techniques such as DamID only reflect single cell organization as a composite or amalgam measure. Another study, employing single cell DamID in a haploid cancer cell line demonstrated that, while there is cell-cell variability, there are core LADs that are consistently maintained at the lamina and that the LADs that do display variability are often developmentally regulated loci (Kind et al., 2015). However, some of these findings are in contrast to numerous studies in primary cells investigating developmentally regulated genes. Many developmentally regulated loci, such as the *Igh* locus, display association with the nuclear lamina, as measured by 3D-FISH, in greater than 90% of cells, suggesting a robust interaction with the nuclear lamina for developmentally regulated genes in relevant cell types (Fuxa et al., 2004; Harr et al., 2015; Kosak et al., 2002; Luperchio et al., 2014; Williams et al., 2006; Wong et al., 2014; Yao et al., 2011; Zink et al., 2004, 2004; Zullo et al., 2012). One caveat to many FISH studies is that they have largely relied on DNA for an individual locus or a discrete number of loci within the nucleus. Such an approach inherently misses some important information—particularly the relationship of all LADs within the chromosome polymer with each other, the lamina, and non-LAD regions. More recently, oligopaint technologies have been employed to identify the disposition of megabase-sized regions or multiple locations within the nuclear volume (Beliveau et al., 2012, 2015; Yao et al., 2011). Since earlier studies using whole chromosome paint techniques suggest that each chromosome occupies distinct territories and self-associated units within the nucleus, we reasoned that examining how LADs organize within an individual chromosome, particularly in relation to non-LAD regions and the nuclear lamina, would offer a more comprehensive view of LAD and overall chromosome organization. While genome-wide molecular methods can be used to predict chromosome and chromosomal sub-domain organization, proof of this functional organization *in situ* requires mapping these interactions back to the single cell.

Here we describe a novel method to directly detect chromosome conformation *in situ* using high-density pools of chemically synthesized oligomers (Oligo Library Synthesis, OLS, Agilent, http://www.genomics.agilent.com/) derived from our DamID data in mouse embryonic fibroblasts (MEFs) (Bickmore, 1999; Kosuri et al., 2010). These high-density oligonucleotides demarcate LADs and non-LAD regions within an entire chromosome, enabling detection of these domains *in situ* and within the context of the entire chromosome polymer. The resulting “chromosome conformation paints” reveal a functional organization of the chromosome territory in single cells heretofore only hinted at by population-based assays such as Hi-C, ChIP and DamID. These chromosome paints identify epigenetic requirements for organization that are missed in high-throughput sequencing-based approaches, and reveal that LADs display two levels of organization: sequestration in the peripheral zone and self-association between LADs on the same chromosome. Integrating these data with Hi-C data in MEFs and pro-B cells, we verify that LADs on a single chromosome self-associate over long distances and that these interactions are related to gene regulation. We also show that LADs comprise the B-compartment and that small interruptions in LADs (Disruption in Peripheral signal, DiPs) correlate with active transcription start sites and enhancers, with a clear switch between compartments (from B to A).

## Results

### LADs in a single chromosome are constrained at the nuclear periphery and peripheral zone

Hi-C (which measures inter- and intra-chromosomal interactions) and DamID (which measures chromatin-lamina interactions) both produce measurements that are an average over large populations of cells. To determine a “preferred” chromosome conformation from these data requires introducing substantial assumptions. Even the best models cannot determine how well the “preferred” conformation is reflected at the single cell level. This can only be measured by direct *in situ* experimental confirmation. To enable such a direct experimental measurement, we derived lamina-chromatin interaction maps in MEFs using DamID (Figure 1A). Specifically, we expressed a chimeric protein comprised of the bacterial adenine methyltransferase (Dam) and the nuclear lamina protein Lamin B1 (LMNB1), to demarcate chromatin that is proximal to the nuclear lamina. Detection was accomplished by deep sequencing. These maps largely agree with those previously published for MEFs (Harr et al., 2015; Figure S1). We next developed a strategy to directly test the compartmentalization of chromosome 11 and 12 in MEFs *in situ*. Previous studies have already shown that each chromosome occupies distinct territories and self-associated units within the nucleus, however, the spatial interactions between individual LADs, the lamina, and the regions between LADs (non-LADs) within a single chromosome have not been investigated. We reasoned that comprehensively examining how LADs organize within an individual chromosome would offer a more comprehensive view of LADs and how they may relate to overall chromosome organization. To design customized chromosome paints for this purpose, we initially identified LAD regions in MEFs using the LADetector algorithm which detects large non-peak like broad domains (https://github.com/thereddylab/LADetector) on previously published DamID data (Harr et al., 2015). The result is a segmentation of the genome into two classes, the LADs and the segments in between them which we designate as “non-LAD”. For each class, 150 base oligonucleotide probes were designed, taking into account GC content, hybridization temperature and filtering for uniqueness of sequence in the genome. The resulting high-density probe pools were comprised of approximately 0.5 million probes per chromosome, divided into LAD and non-LAD pools (Figure 1B; Agilent Technologies). The LAD and non-LAD oligonucleotide pools were chemically coupled to easily distinguishable fluorophores (see Methods) and were hybridized to early pass E13.5-14 primary MEFs using a 3D fluorescence *in situ* hybridization protocol to preserve 3D-organization of nuclei. These two-color probe pools therefore represent a new kind of chromosome paint capable of measuring chromosome domain organization and conformation in single cells.

**Figure 1.**
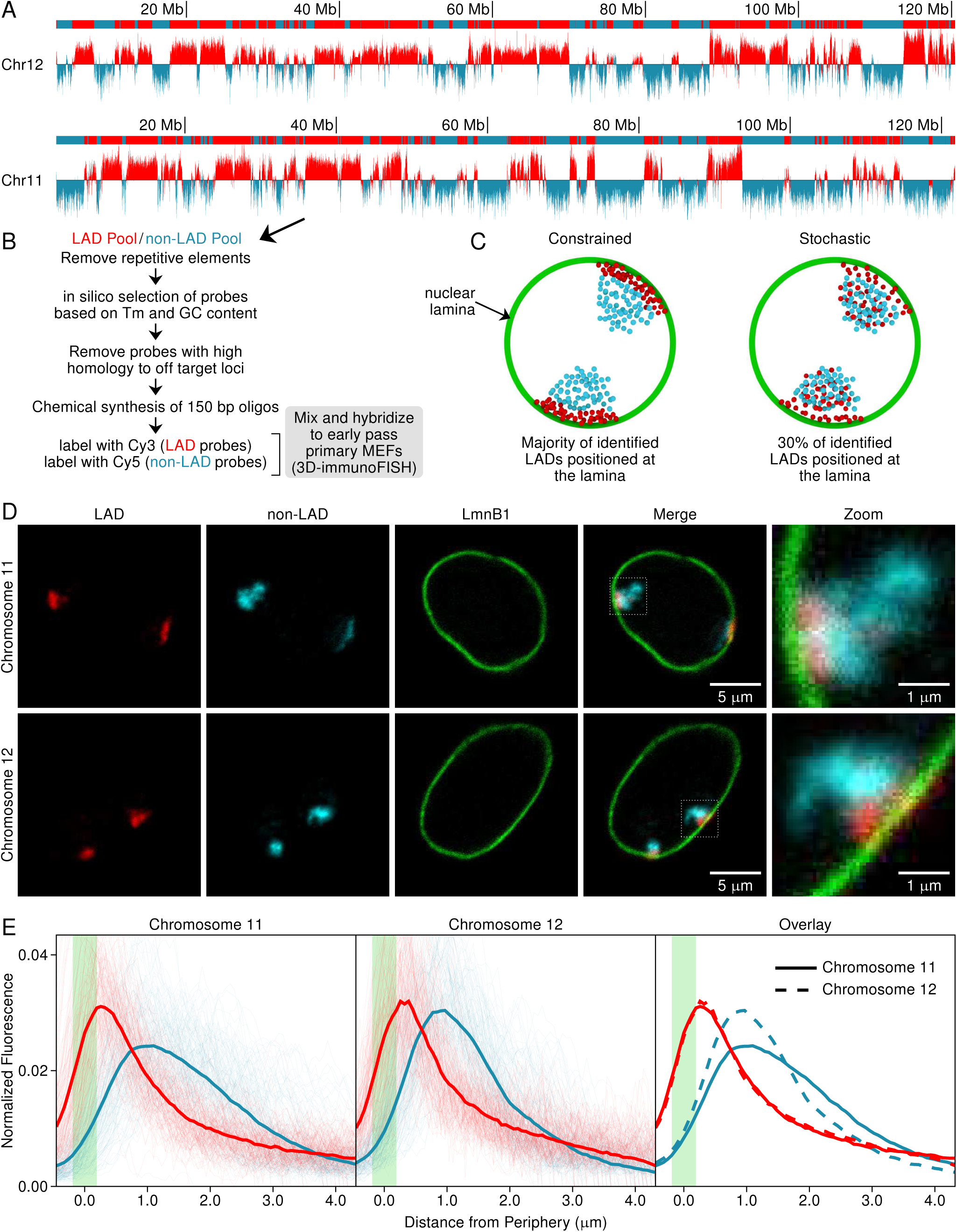
(*facing page*): Chromosome Conformation Paint design and visualization in primary MEFs. (A) LMNB1 DamID log_2_ ratio plots for chromosomes 11 and 12 and LADs (solid red bars) called by LADetector. (B) Workflow for chromosome conformation paint probe design for LAD (red) and non-LAD (cyan) regions. (C)Two hypotheses for the organization of the LAD domains at the single cell level: Constrained (left) in which LADs have anchored or restricted associations with the nuclear periphery and cells exhibit similar organization, or Stochastic (right) in which the population of the cells demonstrate stochastic associations at the nuclear periphery where there is no one uniform organization. (D) 3D-immunoFISH probes in single wildtype MEF nuclei reveals chromosome organization and the presence of LAD and nonLAD subdomains for both chromosome 11 and chromosome 12. (E) Continuous measurements for chromosome 11 (n=50) and 12 (n=50) plotted to show the distributions of the LAD (red) and non-LAD (cyan) signals, as measured from the lamina (green), single cell measurements are shown as thin lines, average as thick lines.

Using these chromosome conformation paints we first set out to directly test whether, *in situ*, the straightforward simple stochastic model holds true, or if, as implied by DamID and developmentally-regulated loci, the organization displays a more constrained configuration (Figure 1C). Using our chromosome conformation paints in a 3D-immunoFISH (FISH on 3D-preserved nuclei with immunofluorescence) assay on early pass primary MEFs, we found clear spatial segregation of the majority of the volume of the LAD and non-LAD domains and their clear organization in interphase nuclei (Figure 1D). To quantify the degree of association with the nuclear lamina, chromosomal subdomains were measured by taking the average of three lines through each territory in the medial planes of the image and plotting the distance from the peak lamin signal for LAD and non-LAD domains (Figure S2). Measurements taken from 50 independent chromosome territories (Figure 1E, Figures S3 and S4) demonstrate that LADs are constrained in close proximity to the nuclear lamina, while non-LAD regions display a broader distribution that peaks nearly a micron or more away from the nuclear lamina (Figure 1E). LADs, in contrast, have a peak signal within 0.25 microns (near the limit of the light microscope) of the peak lamin signal. Moreover, the LAD curves show a striking similarity to one another, even though chromosome 11 and chromosome 12 display different gene densities and LAD composition (Chr11 49.97% LADs and Chr12 62.17 % LADs), indicating that the LADs are forming a structural compartment. The non-LAD curves, by contrast, show no such similarities in structured constraint. Taken together these data show that the LAD sub-domains in a single chromosome are constrained to a peripheral zone at the nuclear lamina.

It is also interesting to note that the LAD and non-LAD compartments occupy not only distinct volumes, but also display differences in their overall volume measurements. 3D image analysis shows that chromosome 11 non-LAD regions occupy an average of 12.3 μm^3^, whereas the LAD sub-territory occupies 7.71 μm^3^. In contrast, LADs on chromosome 12 occupy about 4.3 μm^3^ and non-LADs occupy 7.8 μm^3^. Interestingly, chromosome 12 (overall) is more compacted than chromosome 11, even though the two chromosomes are nearly identical in overall length. This is at least partly explained by the fact that the LAD regions on chromosome 12 cover a greater percentage of the chromosome (see Figure 1A). These data strongly suggest that the LAD sub-territory compartment is “more compact” than the non-LAD, in agreement with previous studies. Strikingly, even though there is a disparity in the *overall* volume of the LADsub-territories between chromosome 11 and 12, the distance that they radiate from the nuclear periphery is nearly identical, suggesting that constraint at the nuclear lamina results from the physical properties of that interface and not from overall LAD density. This region where LADs interface with the edge of the nucleus and are restricted we therefore describe as the “peripheral zone”.

### LADs are the B-compartment

Given the data above, which seemed to indicate that LADs are constrained into a single nuclear sub-compartment in proximity to the nuclear lamina, we next asked how these domains related to the chromatin A/B-compartments, which are bioinformatically derived from Hi-C data. Hi-C comprehensively detects chromatin interactions across the genome. This method is based on 3C, in which chromatin is crosslinked with formaldehyde, digested, and then re-ligated so that only DNA fragments that are covalently linked together form ligation products. The resulting ligation products are then subjected to massively parallel paired-end deep sequencing. The sequences are mapped back to the genome and interactions between chromosomal segments in 3D are captured. These data can be used to detect more local self-interacting domains (TADs) or to determine domain interactions across the entire chromosome (A/B-compartments). A Hi-C derived “compartment” is comprised of regions of the genome that interact with each other. In these data, regions in the A-compartment display preferential interaction with other regions in the A-compartment, and regions in the B-compartment with other regions in the B-compartment. The A-compartment is thought to harbor more “active” regions of the genome, while the B-compartment is thought to be more “repressive”.

While previous studies have shown an enrichment of LADs in the B-compartment through bulk correlation analyses, a direct comparison between the domain structures has not been undertaken (Rao et al., 2014; Schwarzer et al., 2016). To enable such a direct comparison, we used publicly available Hi-C data in MEFs (Krijger et al., 2016) and a new high-resolution compartment-calling approach to investigate the relationship between LADs and the B-compartment (Sauria et al., 2015). Prior compartment calling approaches have used inter-chromosomal data and relied on lower-resolution binning to overcome the sparseness of longer-range Hi-C interaction data in creating interaction correlation matrices and finding the first eigenvector to determine compartments (Lieberman-Aiden et al., 2009; Rao et al., 2014). We previously demonstrated that by using a dynamic binning-based approach, inferring interaction enrichments over differing ranges based on data sparseness coupled with eigenvector decomposition we could achieve a significantly higher resolution partitioning of the genome into chromatin compartments (Jung et al., 2017). To further improve this, we developed a likelihood-based compartment score. Counts were assumed to be distributed under the Poisson distribution with differing decay rates with distance for interactions within or between compartments:

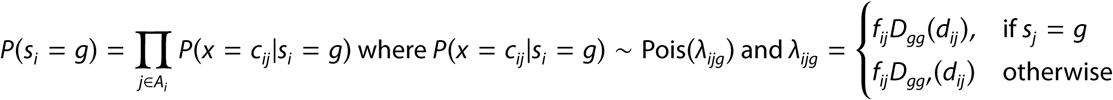
where *c*_*ij*_ is the read count between fragments *i* and *j, s*_*i*_ is the compartment state of fragment *i, f*_*ij*_ is the sum of interaction normalization values for bin *ij, d*_*ij*_ is the inter-fragment distance, and *D*_*gg*_ is the distance decay function for interactions both in state g. The compartment score is then Score (*i*) = log_2_ (*P* (*s*_*i*_ = *B*)/ *P* (*s*_*i*_ = *A*)) or the log ratio of the relative probability of occurring in the B-compartment vs. the A-compartment. Using the high-resolution eigenvector-based compartment calls, we initialized the compartment states for the model at a resolution of 10 kb and trained using an expectation-maximization approach. Compartment calls were highly consistent with both low and high-resolution eigenvector-based methods (Figure S5) although compartment boundaries and small-scale features were refined. This approach marked an improvement over both previous approaches in that it maintained a higher-resolution partitioning without needing to use neighboring data (thus smoothing over features) for data-sparse regions. Using our high-resolution compartment scoring method, we observed a striking similarity between compartment scores and LAD DamID scores in MEFs (Figure 2A and 2B). The B-compartment and LADs not only share the same genomic regions but also have nearly identical boundaries (Figure 2A and 2C). While DamID biological replicates show an agreement of around 91%, more than 80% of the genome in MEFs share consistent state calls (B-compartment/LAD or A-compartment/non-LAD) across these orthogonal measures. The regions of disagreement are highly enriched in ambiguous compartment scores (close to zero), representing regions of low Hi-C sequencing coverage, transient lamina association, or mixed state across the population. For example, the well-studied 3 Mb *Igh* locus, which itself occupies an entire LAD in MEFs, is also contained in a B-compartment domain nearly identical to the corresponding LAD (Figure 2C). We do note that the compartment and DamID scores are not identical, although the clear majority of differences share a common feature. DamID identifies sequences that were marked as lamina proximal at any point during the assay, regions that are transiently or stochastically associated with the lamina are labeled while Hi-C represents a snapshot of a single set of conformations. As such, transient or dynamic chromatin arrangements show much less robust compartment preferences.

**Figure 2:**
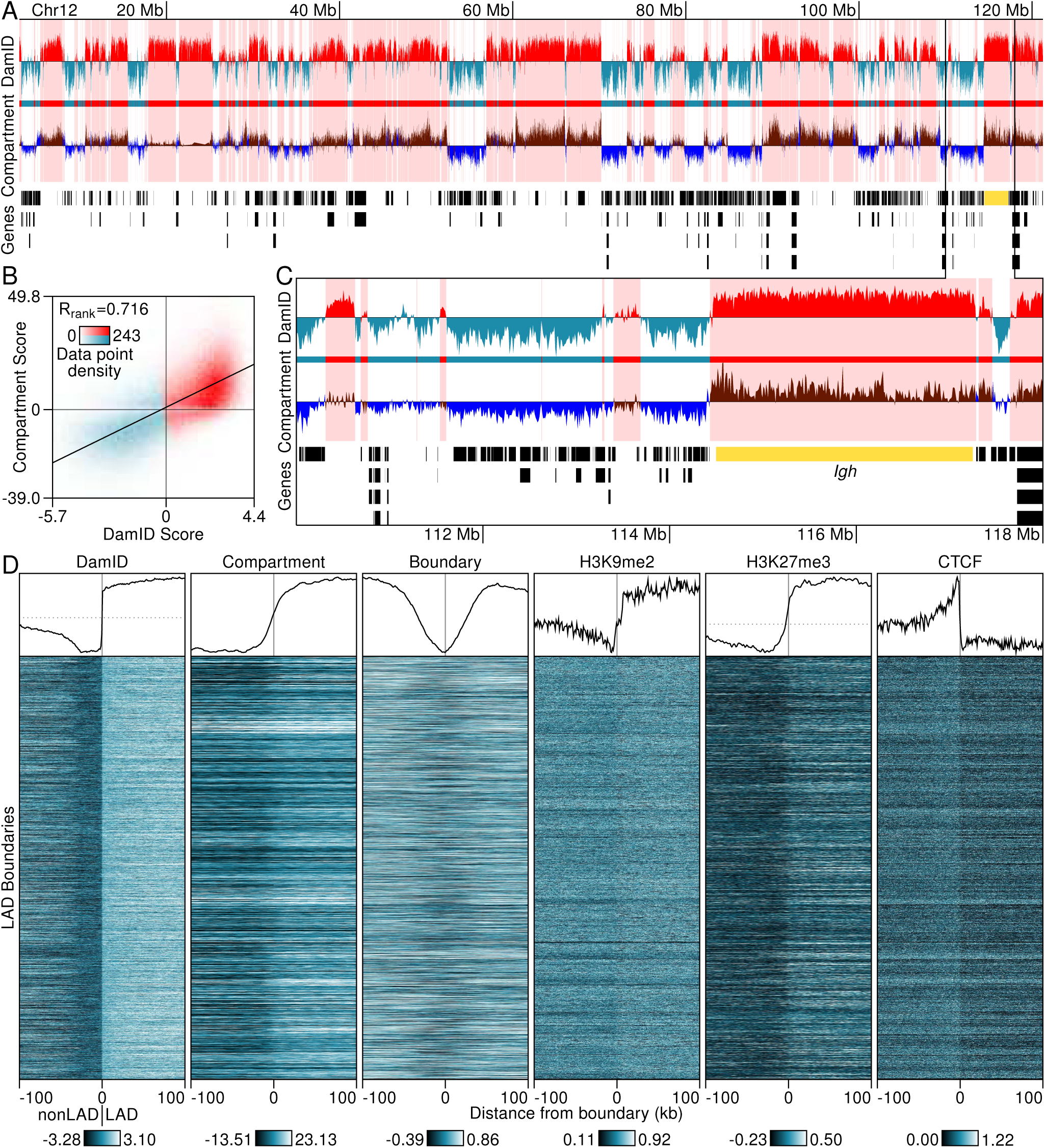
LAD structures captured by both local and chromosome-wide metrics from Hi-C data. (A) LMNB1 DamID, Hi-C compartment score, and UCSC genes from chromosome 12 in MEF cells. LAD calls and associated data are highlighted in red/brown (DamID/compartment scores) while data from non-LAD regions are shown in blue/purple. (B) Genome-wide correlation between DamID and compartment scores. Data are partitioned into a 100 by 100 grid with intensity indicating data density and color showing whether the majority of the bins data points are in LADs (red) or not (blue). (C) Zoomed view of the *Igh* locus (yellow) and the 8 Mb upstream. (D) Feature profiles anchored at all boundaries of LADs of size 100 kb or greater (excluding chromosome X) and oriented from nonLAD (left) to LAD (right). Line graphs (top) represent the average trend across all boundary profiles for each feature. Profiles consist of data within 100 kb of each boundary binned in 1 kb intervals. Bins with no data are shown in gray.

Given these data, we sought to determine the extent to which LAD/non-LAD and A/B-compartment boundaries aligned (Figure 2D). Specifically, we identified all LAD/non-LAD boundaries (+/-100 kb, Figure 2D DamID) and, using these genomic intervals, aligned the A/B-boundaries relative to the LADs. Using this approach, we note a striking concordance of LAD and A/B-compartment boundaries (Figure 2D, compartment). In addition, we computed a Boundary Index score, which is a local measure of how frequently interactions occur between regions upstream and downstream of a given site (250 kb) in the Hi-C data (Jung et al., 2017). This additional boundary measure also aligns with LAD boundaries (Figure 2D, boundary). As has previously been noted, LADs are enriched for H3K9me2/3 and H3K27me3, and in our analyses both LADs and the B-compartments are enriched in these modifications. As a further indication that the LAD and compartment boundaries demarcate two distinct functional domains, the chromatin organizer CTCF is enriched at the B-compartment and LAD boundaries, and is depleted within LADs and the B-compartment.

### LAD organization is dependent on both chromatin state and A-type lamins

Previous studies have demonstrated that the organization of at least some LADs to the nuclear lamina is dependent upon H3K9me2/3 and H3K27me3, and that the accumulation of these histone modifications may contribute mechanistically to LAD formation and maintenance. We therefore next asked if disruption of heterochromatin through drugs targeting specific epigenetic modifiers affects LAD organization. Specifically, we treated with Trichostatin A (TSA, an HDAC inhibitor that promotes histone acetylation), BIX01294 (which inhibits H3K9me2 through inhibition of G9a and G9a-like protein) or 3-Deazaneplanocin A (DZNep, which decreases H3K27me3 through inhibiting EZH2). We have previously shown that loci in LADs are able to be redirected away from the lamina upon disruption of either H3K9me2/3 or H3K27me3. To test whether LADs are generally disrupted by these treatments, we subjected pre-treated cells to the DamID protocol to ensure that we were measuring the reconfigured organization of LADs. Surprisingly there was little or no difference in LAD organization by these measures (Figure 3A). Disruptions using DZNep and BIX01294 displayed 93% to 96% overlap of LAD organization with non-treated cells. TSA treatment showed modestly more derangement, with 86% overlap of LADs between non-treated and treated cells. Intriguingly, LAD disruption in TSA conditions appears to occur over regions that displayed low lamina signal in untreated cells, suggesting that these regions are more susceptible to disruption.

**Figure 3:**
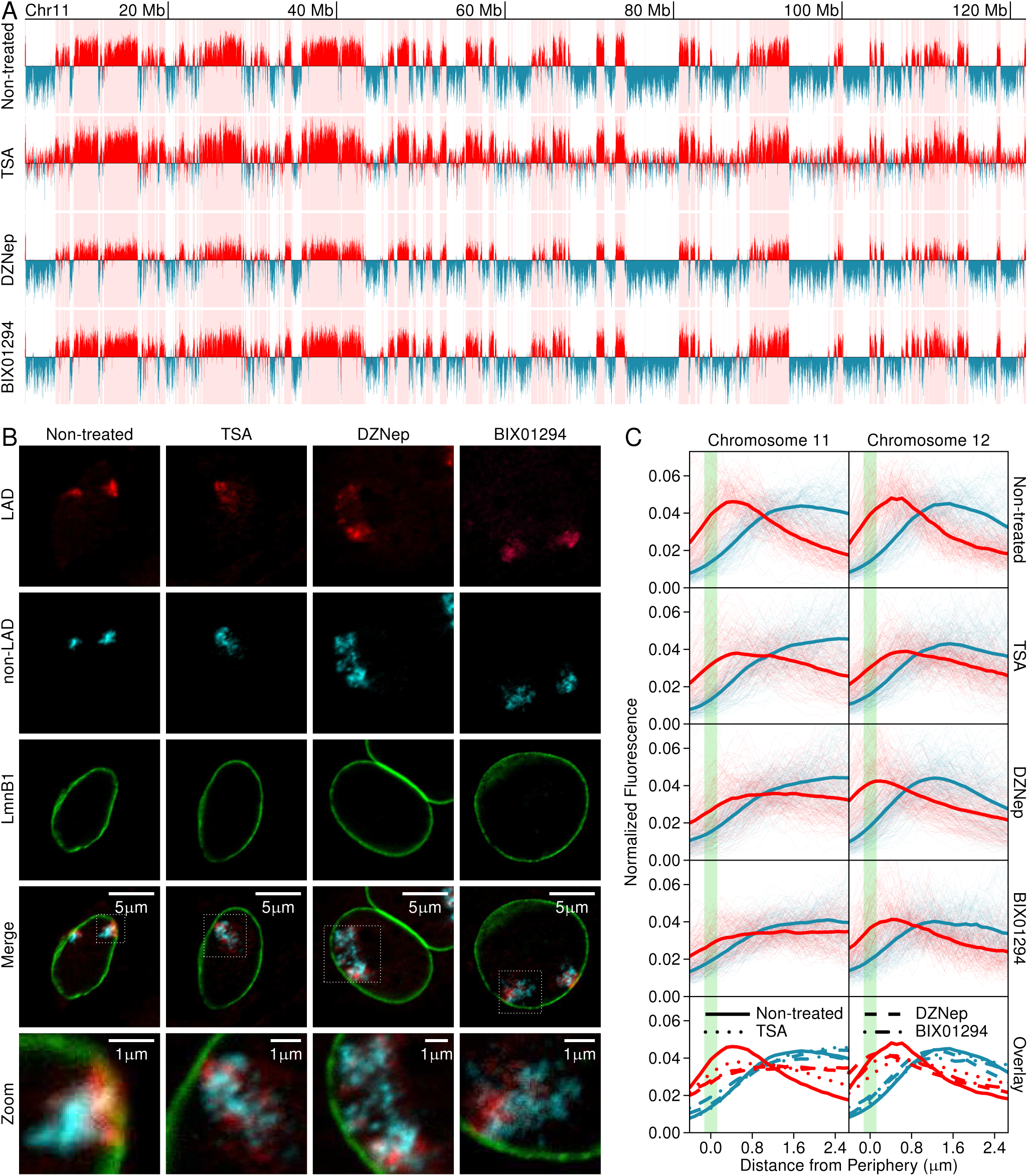
Epigenetic Perturbation and Sub-chromosomal Architecture. (A) DamID measurements for non-treated and drug-treated MEFs. (B) 3D immunoFISH signals of LADs and non-LADs are visualized in non-treated and drug treated nuclei. (C) Individual measurements show distributions of LAD (red) and non-LADs (blue) relative to Lamin B1 (green). Individual measurements of chromosome territories are shown as thin lines. Overlays of distributions for chromosomes 11 and 12 are provided in bottom two graphs. Non-treated Chr11 n=50, Chr12 n=50; TSA treated Chr11 n=52, Chr12 n=51; DZNep treated Chr11 n=52, Chr12 n=54; BIX01294 treated Chr11 n=51, Chr12 n=50.

Our previous studies suggest that both an ectopically lamina-targeted locus and endogenous loci moved away from the nuclear lamina upon perturbation of either H3K27me3 or H3K9me2/3, in contrast to what is observed by DamID (Figure 3A). We hypothesized that the DamID assay, an ensemble measure of lamin association across a population of cells, may be smoothing out the level of disruption displayed at the single cell level. To test this, we performed 3D-immunoFISH using our chromosome conformation paints to detect, at the single cell level, the relative disposition of the LAD and non-LAD chromosome nuclear sub-compartments to each other and the nuclear lamina (Figure 3B; Figures S6 and S7). The level of disruption of LAD organization in individual cells was quite striking and varied from cell to cell. Many territories were so completely disrupted by perturbations that they were unable to be scored by our methodology as we were unable to identify distinct chromosome territories or identify medial plane measurements. In addition, these treatments altered the morphology of the nuclei. Nonetheless, we are able to detect substantial reorganization, even in the cells with intact chromosome territories. To provide consistent measurements for all conditions, we identified territories in which the majority LAD signals were in the medial planes. We note that this approach often led to selection of nuclei with less perturbed chromosome organization. In all conditions some lamina contacts remain, perhaps explaining in part the DamID data. However, many LADs relocate away from the lamina and, notably, one of the most obvious disruptions is the loss of the self-association of the LADs along with a general expansion of the sub-territory. This disorganization is not simply a consequence of the altered nuclear morphology, but rather shows intermingling of LADs and non-LADs, a general loss of LAD-to-LAD interactions, and loss of distinct compartmentalization (Figures S6 and S7). Treatment with TSA resulted in the most obvious disruptions of the territories, in both loss of self-association and disassociation from the lamina, which may also be reflected in the DamID scores, with a loss of clear distinction between LADs and non-LADs. We observe less of an effect on chromosome 12, which is more compact (and LAD-rich) and may display a greater resistance to perturbations in the time-frame tested (24-48 hour treatment). Overall, notwithstanding the population measures, these data demonstrate that the LAD compartment is comprised of two types of organization, LAD-to-LAD interactions and restriction to the peripheral zone, and both are dependent upon chromatin state. We note that the loss of LAD-to-LAD interactions have been missed by other imaging-based approaches, demonstrating the utility of our chromosome conformation paints. These data also demonstrate a significant amount of decompaction of the total chromosome territory (LAD and non-LAD), consistent with previous studies.

Previous studies have also implicated the role of Lamin A/C in regulating LAD organization at the nuclear lamina. Lamin A/C has been shown to be required to tether heterochromatic regions to the nuclear periphery in differentiated cells (Solovei et al., 2013). More recently, studies in embryonic stem (ES) cells have shown that the lamins may be dispensable for organizing LADs, although another study re-examining those data suggest that LAD organization is affected (Amendola and van Steensel, 2015; Zheng et al., 2015). Our previous data in MEFs suggests that, at least for the loci tested by FISH, Lamin A/C is required for maintenance and/or establishment of LADs in these cells (Harr et al., 2015). In order to test the role that Lamin A/C plays in organizing LADs, we removed Lamin A/C by shRNA-mediated knockdown. Knockdown with this construct is nearly complete and does not affect either H3K9me2/3 or H3K27me3 (Figure S8A). Surprisingly, even complete knockdown of Lamin A/C does not result in an altered LAD profile by DamID (Figure 4A), in agreement with previous studies on ES cells. We next asked if we could detect any alterations in LAD organization (either association with the lamina or the self-association of LADs) using our chromosome conformation paints. We note that the DamID and *in situ* hybridization studies were done in parallel, taking cells from identically treated cell culture plates. The cells to be used for 3D-immunoFISH were taken at the time when the lentiviral DamID constructs were added to the culture dishes, ensuring that both methods are measuring the same level of disruption. In contrast to the DamID assays, which measure populations of cells over time, the 3D-immunoFISH assays with the chromosome conformation paints uncover a much more disorganized chromosome organization at the single cell/single time point level (Figure 4B and 4C; Figure S8B and S8C).The disruptions to normal LAD and non-LAD organization were so extreme that many nuclei had to be dropped out of the analyses, as we could not discriminate chromosome territories. These disruptions included severe decompaction of the chromosome territory, intermingling of LAD and non-LAD signals, loss of peripheral association of LADs and loss of LAD-to-LAD aggregation. In addition, as noted for the drug treatments, the overall nuclear morphology changes upon Lamin A/C knockdown, potentially confounding some of our results. Nonetheless, it is obvious from these assays that the A-type lamins are critical for organizing LADs at the lamina, and for either establishing or maintaining self-association of LADs.

**Figure 4:**
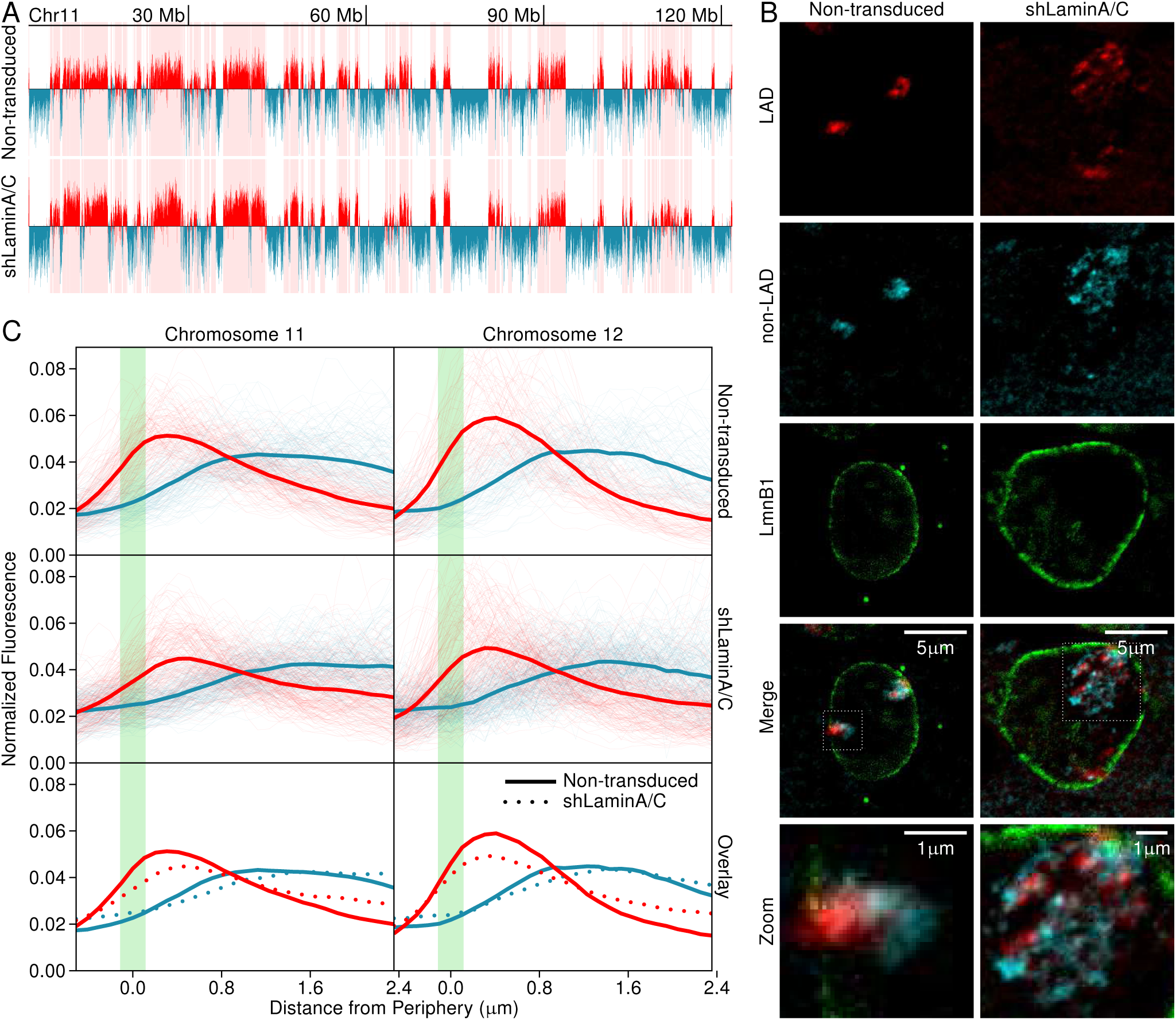
Nuclear Structure Integrity and sub-chromosomal Architecture. (A) LMNB1 DamID data for Wildtype vs LaminA/C knockdown for chromosome 11. (B) 3D immunoFISH signals of LADs and non-LADs in wildtype and LaminA/C knockdown highlight chromosome sub-territory organization after LaminA/C knockdown. (C) Composite measurements for chromosomes 11 and 12. Individual measurements are shown as thin lines and averages as thick lines. Wildtype Chr11 n=44, Chr12 n=22) and LaminA/C (Chr11 n=60, Chr12 n=66)

### Developmental reorganization of chromosome conformation

Having demonstrated the dependence of LAD organization on chromatin state and A-type lamins using these perturbation strategies, we next sought to investigate reorganization of LADs in a normal biological state. To link these observations to a significant developmental process, we considered a key developmentally regulated region, the *Igh* locus, which occupies approximately 3 Mb of the distal half of chromosome 12 in Murine cells (Figure 5A).The *Igh* locus is lamina proximal in non-B lineage cells, when the locus is transcriptionally inactive, and has chromatin modifications consistent with a repressed state (Bertolino et al., 2005; Kosak et al., 2002; Reddy et al., 2008). During development and specification to the B cell lineage (at the pro-B cell stage) *Igh* moves away from the lamina and displays loss of H3K9me2/3 and H3K27me3 concomitant with its activation (Bertolino et al., 2005; Fuxa et al., 2004; Johnson et al., 2003, 2004; Reddy et al., 2008). We next used the chromosome conformation paints for chromosome 12 (designed to the LAD organization in fibroblasts) to determine if they could detect the reorganization of the 3 Mb region in pro-B cells. We first mapped the LAD organization in pro-B cells and fibroblasts using DamID (Figure 5A). As we previously noted, LAD organization in pro-B cells and fibroblasts is largely unchanged (Harr et al., 2015; Zullo et al., 2012). However, key regions do reorganize between the cell types, including the *Igh* locus. It is important to note that, unlike the pleiotropic drug treatments where it was difficult to determine specific regions that changed by DamID (Figure 3A), this type of developmentally directed reorganization led to very clear, robust and domain specific reorganization that was easily revealed by DamID (Figure 5A), suggesting that this region is no longer in proximity with the nuclear lamina in the majority of the cell population. This is consistent with existing 3D-immunoFISH studies of the locus (Kosak et al., 2002; Reddy et al., 2008). To test the interaction of the *Igh* locus with the lamina or LAD/non-LAD compartments in single cells using our chromosome conformation paints, we employed a four-color 3D-immunoFISH strategy demarcating the lamina (green), LADs (red), non-LADs (cyan) and the *Igh* locus (yellow, Figure 5B and 5C). As expected, in MEFs the *Igh* locus is lamina proximal and located within the LAD compartment. In contrast, in pro-B cells the *Igh* locus has escaped the peripheral zone and is no longer associated with the other LADs that are still found proximal to the nuclear lamina. We note that the probe used for the *Igh* locus is approximately 150 kb in size (CT7 526A21), while the locus itself is 3 Mb. In addition, a region that is configured in a LAD in MEFs adjacent to the *Igh* locus is also reconfigured away from the lamina in pro-B cells (Figure 5A). Thus, the *Igh* specific probe (yellow) only marks a small portion of this larger reorganized domain. These data suggest that developmentally driven reorganization of loci can move LADs away from the nuclear lamina and abrogate self-interaction between these regions and regions that remain in a LAD.

**Figure 5:**
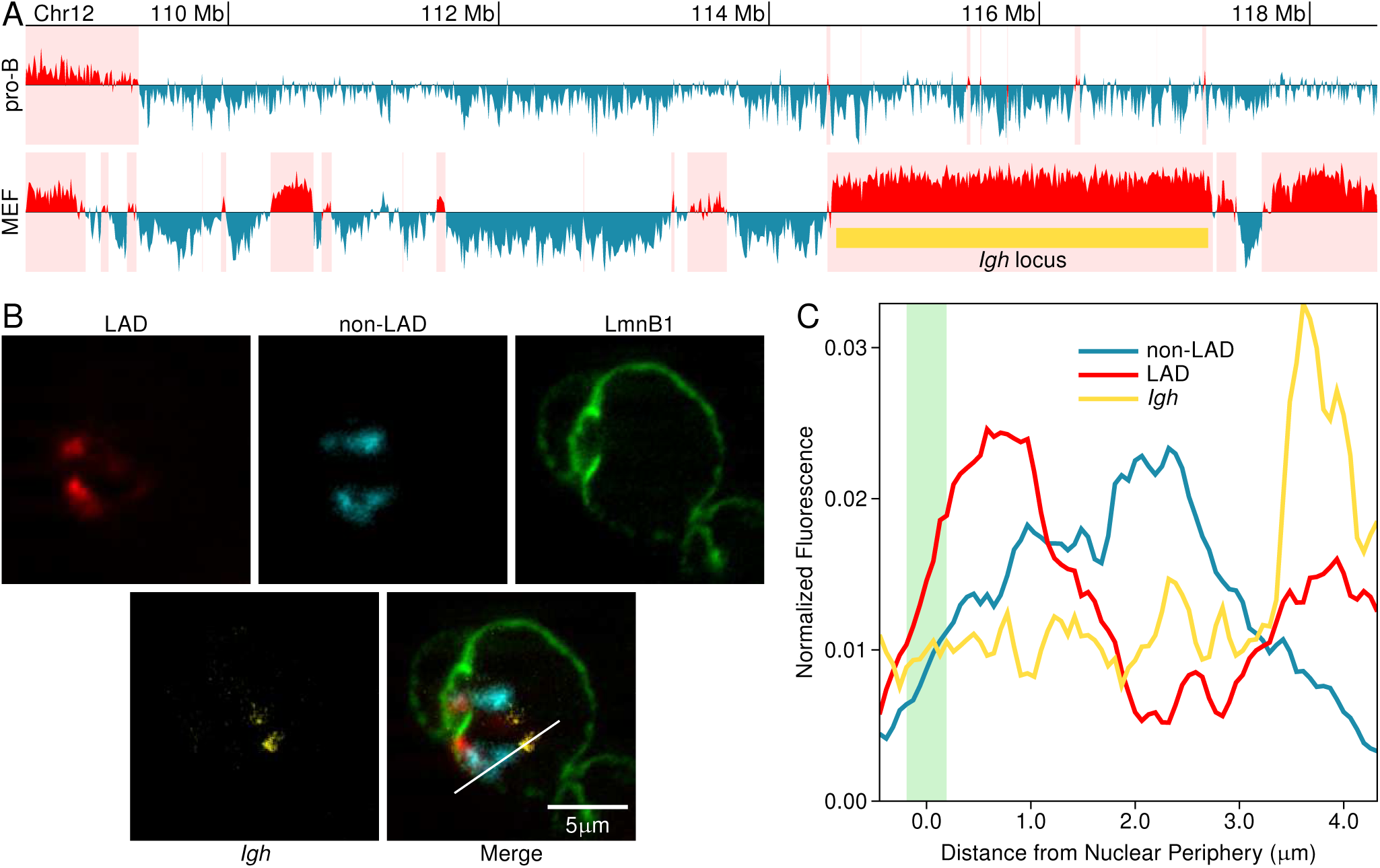
Subchromosomal reorganization in a developmental model. (A) LMNB1 DamID data in pro-B and fibroblast cells show the *Igh* locus (yellow bar) is in a vLAD. (B) 3D-immunoFISH with probes designed in MEFs show organization of LADs (red) and non-LADs (cyan) in pro-B cells. The *Igh* locus is centrally disposed in pro-B cells and in a vLAD (*Igh* in yellow, panel B). (C) Distributions of LAD, non-LAD and *Igh* locus relative to Lamin B1 along the white line indicated in panel B merge image.

Having shown that LADs self-interact using imaging, we next determined if these long-range associations could be detected by Hi-C and A/B-compartment analysis. Since the *Igh* locus shows robust reorganization both by population measures (DamID) as well as at the single cell level (3D-immunoFISH), we used publicly available Hi-C data to compare chromosome organization in MEFs and pro-B cells with the *Igh* locus as an anchor point for comparing compartmentalization and interactions with other LADs. We first measured the degree to which LADs identified in MEFs interacted with either each other or non-LAD regions (Figure 6A). LAD regions do indeed show an increased interaction with other LADs, and non-LAD regions also self-interact. This is consistent with our previous data demonstrating that LADs represent the B-compartment (Figure 2) and the B-compartment itself is a measure of self-interactions. As the *Igh* locus undergoes large-scale reconfiguration away from the lamina between MEFs and pro-B cells, we next queried whether these domains also lost their long-range associations with other LADs once they moved away from the nuclear lamina, and presumably, out of the B-compartment. We generated heat maps representing levels of interaction between all regions on chromosome 12 (Figure 6B).The Hi-C data clearly support the imaging data suggesting that LADs self-interact over large distances. Focusing on the *Igh* locus, we mapped interactions of this domain and adjacent compartments from Hi-C data in both MEFs and pro-B cells (Figure 6C). Remarkably, the *Igh* locus (highlighted in yellow) changes its interaction with the rest of the chromosome in pro-B cells compared with MEFs, while most LADs do not change substantially between cell types (red bars). In MEFs, when the *Igh* locus is in a LAD, the interactions of this region with the rest of the chromosome are almost exclusively with other LADs. Interestingly, in MEFs the LAD harboring the *Igh* locus does not have many interactions within its own domain, nor does it interact with an adjacent LAD, suggesting some local constraint preventing their interaction. In contrast, in pro-B cells, this region displays robust self-interaction within the domain (Figure 6C, lower panel). These findings are supported by previous 3D-immunoFISH studies which show this locus in a more extended conformation in non-B cells compared to pro-B cells. Even more strikingly, the compartment encompassing the *Igh* locus switches its affinity away from LADs and now interacts almost exclusively with non-LAD regions.

**Figure 6:**
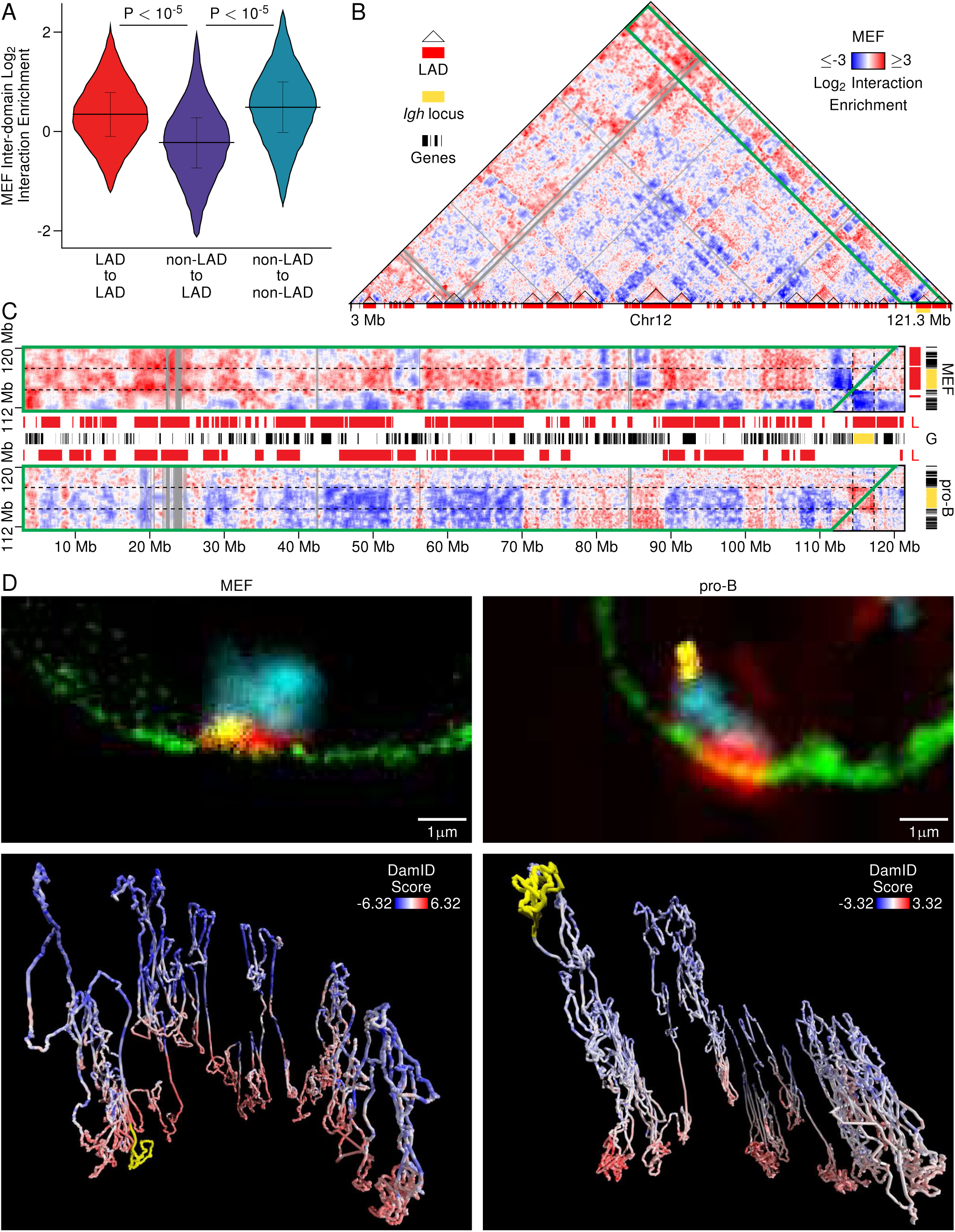
(*facing page*): Associations between LADs are seen across whole chromosomes and in a cell type-dependent manner. (A) Violin plot depicting interaction enrichments across all pairs of LAD and/or non-LAD regions in MEF cells. P-values correspond to both the Mann-Whitney U test and the independent sample t-test comparing within LAD/within nonLAD interactions versus between LAD and nonLAD interactions. (B) Dynamically-binned 50 kb enrichment heatmap of chromosome 12 MEF Hi-C data. All bins are expanded to contain a minimum of 10 reads. Regions with no data (filtered out) are in gray. LADs are outlined in black and the *Igh* locus is marked by a yellow rectangle. (C) A slice of dynamically-binned enrichment interaction data from chromosome 12 spanning the whole chromosome along the X axis and a window centered on the *Igh* locus along the Y axis. The position of the *Igh* locus-containing LAD from MEF cells is marked with dashed lines. The letters ‘L’ and ‘G’ mark the LAD and gene tracks, respectively. (D) Single cell 3D-immunoFISH probe images depicting chromosome 12 LADs (red), nonLADs (blue), LmnB1 (green), and the *Igh* locus (yellow) for MEF and Pro-B cells (top) and Hi-C consensus models for these cell types colored by cell type specific DamID scores and with the *Igh* locus identified in yellow (bottom).

To visually compare the conformation implied by Hi-C to that observed with chromosome conformation paints, we constructed 3D models of chromosome 12 by performing principal component analysis on pairwise distances inferred from MEF and pro-B Hi-C data, and compared them to corresponding 3D-immunoFISH images (Figure 6D). There is a striking correspondence between the computational models and single cell images, again showing that, while most LAD regions do not change their localization between MEF and pro-B cells, the *Igh* locus undergoes dramatic and visible reorganization within the chromosome. Overall the domains surrounding the *Igh* locus change their interactions concomitant with their organization relative to the nuclear lamina. In addition, the changing LAD landscape correlates with a changing chromosome conformation across the entirety of the chromosome (Figure S9). Taken together, these data strongly suggest that LAD organization is tightly coupled with overall 3D organization and chromosome conformation. These data also highlight that the ensemble measures of HiC and DamID, at least on a global level, are reflective of chromosome architecture in single cells.

### Sub-structure of LADs and B-compartments reveal potential regulatory elements

Lamina associated domains comprise large (>100 kb) genomic regions. However, within these domains exist smaller regions that have low or negative enrichment for interactions with the nuclear lamina, which we have dubbed “DiPs” (Disruption in Peripheral signal). Similar breaks in otherwise large contiguous domains have previously been observed for H3K9me2 (Wen et al., 2012). We defined DiPs as regions of 1-25 kb in LADs that display negative interaction with the nuclear lamina by DamID. We identified all DiPs present in MEFs and evaluated the distribution of various epigenomic features in a +/-100 kb region around each (Figure 7). The DamID signal distribution show that these were indeed discrete regions of lamina low signal (Figure 7, DamID). We noticed that DiPs contain transcription start sites (TSSs) at a frequency more than twice what would be expected (p<0.001), although the majority of these DiPs do not harbor an annotated TSS. For graphical clarity, we have subdivided the DiPs into those with at least one TSS present (Figure 7, gold) and those with no annotated TSS (Figure 7, blue). We next asked if these small interruptions in LAD organization correlated with changes in A/B-compartment assignment or were enriched for boundary features by Hi-C data, as was done for Figure 3. We observe a general correlation of these regions with a shift in compartment association, especially for DiPs containing at least one TSS, an observation made possible by our finer scale likelihood-based scoring approach. We also observe a measurable boundary signature for TSS-containing DiPs (Figure 7, Compartment and Boundary columns). One discrete measure of boundary potential is the occupancy of either CTCF or cohesin within these regions(Dixon et al., 2012; Nora et al., 2012; Phillips-Cremins et al., 2013; Rao et al., 2014; Sexton et al., 2012). Both CTCF and cohesin showed enrichment in DiPs within LADs, irrespective of whether these regions harbor a TSS, supporting the boundary analysis that these regions demarcate a structural shift from one compartment to another. H3K27me3 has been shown to be enriched in LADs, particularly at border regions. Strikingly, only a subset of these DiPs displayed H3K27me3 enrichment, suggesting that even though these regions are behaving like structural borders, they are not behaving like LAD borders (Figure 7, H3K27me3). Intriguingly, the DiPs, irrespective of TSS status, were enriched in H3K27ac, indicating that these regions represented a more euchromatic environment than the adjacent LAD regions (Figure 7, H3K27ac).

**Figure 7:**
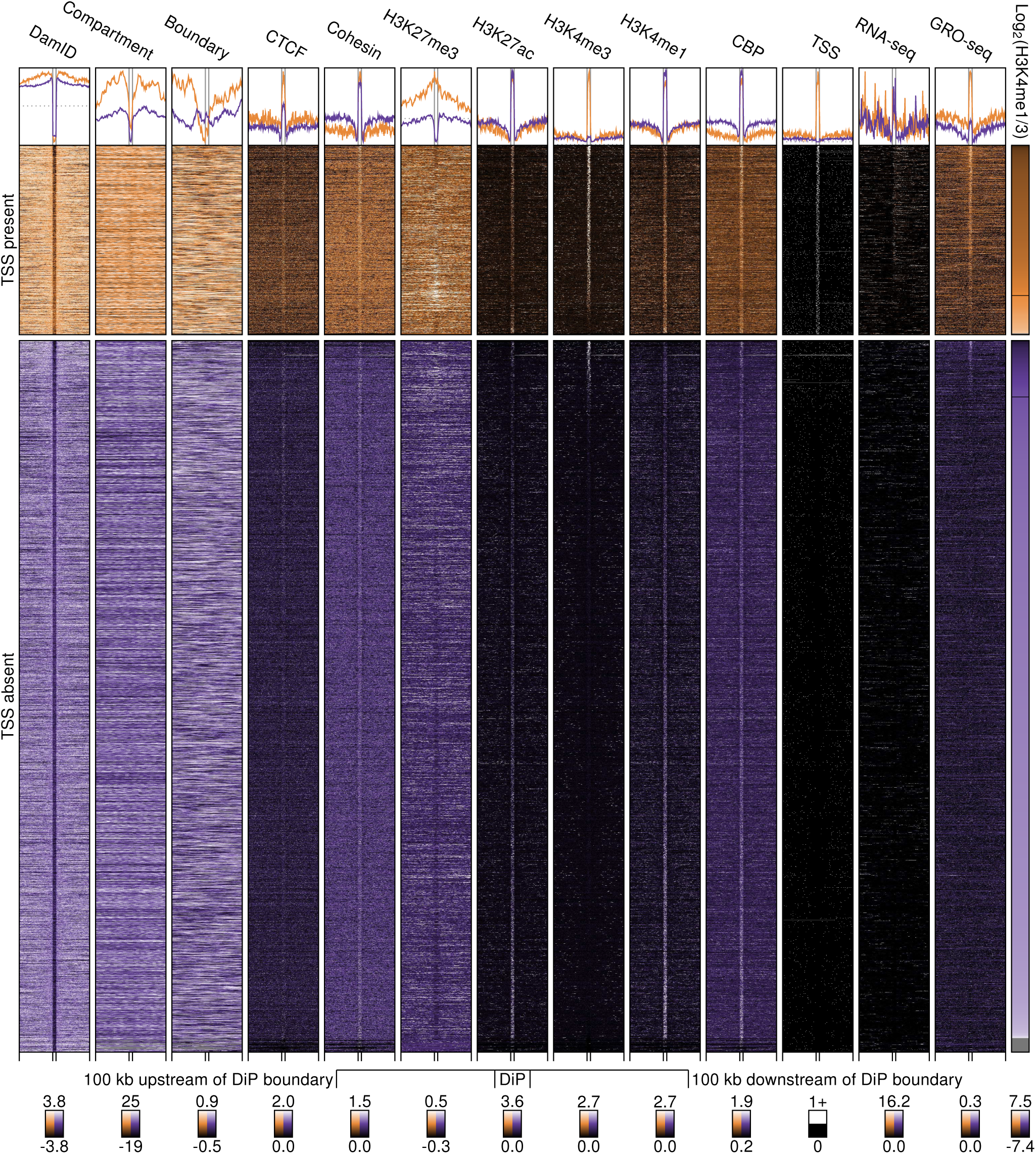
LAD DiPS features reveal potential regulatory role. Feature profiles are shown for all DiP calls in MEF cells (excluding chromosome X), extending 100 kb away from DiP boundaries. Feature data are binned at 1 kb intervals up- and downstream of DiPs and in 10 evenly sized intervals across each DiP. Profiles are partitioned into those containing at least one TSS (from UCSC gene set) within the DiP boundaries (orange) and those that do not (purple). The right column shows the ratio of H3K4me1 to H3K4me3 mean signals within the each DiP. Columns are sorted from lowest to highest ratio. Line graphs (top) represent the average trend across all boundary profiles for each feature.

Given these data, we next queried whether some of these regions might represent either active TSSs, poised TSSs, or enhancers. In order to determine if a DiP harbored an active TSS, we compared CBP occupancy, GRO-seq data, steady-state RNA levels, and H3K4me1/me3 levels (Barski et al., 2007). A majority of the TSS-harboring DiPs display characteristics of an active transcription start site including production of RNA, occupancy by CBP, and a low H3K4me1/H3K4me3 relative ratio (Figure 7). However, there are a subset of these TSSs that display a low H3K4me1/H3K3me3 ratio and low or no RNA transcription even though occupied by CBP. These TSSs appeared to be in a “poised” state (Figure 7). A few of the DiPs containing no TSSs showed hallmarks of an active promoter region, and these could represent previously unannotated transcription start sites. The majority of the DiPs, however, appear to represent enhancers as determined by a higher H3K4me1/H3K4me3 relative ratio (DiP enhancers). We note that the adjacent LAD intervals do not show enrichment for boundary markers, active transcription or putative enhancer activity. It is important to note that these DiPs are associated with a dramatic change in A/B-compartment assignment, suggesting that regions localized to A and B-compartments can be much smaller than previously noted. In addition, unlike the LADs, these DiPs preferentially interact within the A-compartment, suggesting that these regions must loop out of the LAD/B-compartment.

One question that arises from the previous data is whether DiP enhancers display a clear interaction with other regulatory elements in the A-compartment. We used the MEF Hi-C data to determine if we could detect specific interactions of a DiP enhancer with other regulatory elements in the genome. The heatmaps in Figure 8A and 8B show two examples of putative enhancers that show preferential interaction with the promoter region of a gene adjacent to, but outside of, the LAD. These interactions can also be detected by focusing on the enhancer (enhancer-anchored) and measuring (in sliding 1 kb bins) its interaction with these more distal regions, or by focusing instead on the promoter region and looking for preferential interactions with any region inside the LAD using a sliding 1 kb window. In both cases a preferential and specific interaction is noted between the enhancer located in a LAD-encompassed DiP and the promoter located outside of the LAD. We note that the majority of enhancers and promoters encompassed in LADs are not in DiPs, and that even if an enhancer or promoter is looped out from a LAD in a DiP it is not always possible to detect specific interactions by Hi-C. As has previously been shown, LADs harbor relatively inactive genes. By comparing the relative steady-state transcript levels between LADs and non-LADs, this trend is obvious (Figure 8C and 8D). However, we note that there are exceptions to this rule, and we propose that these regions within a LAD that display higher levels of transcription represent regions that have looped away from the lamina, as evidenced by low signal by DamID and their increased interactions with the A-compartment. While the majority of transcription start sites in LADs are inactive and display high levels of interaction with the lamina as determined by DamID score, a significant proportion appear to lose their tight association with the lamina and this is concordant with a relative increase in transcriptional activity for this population.

**Figure 8:**
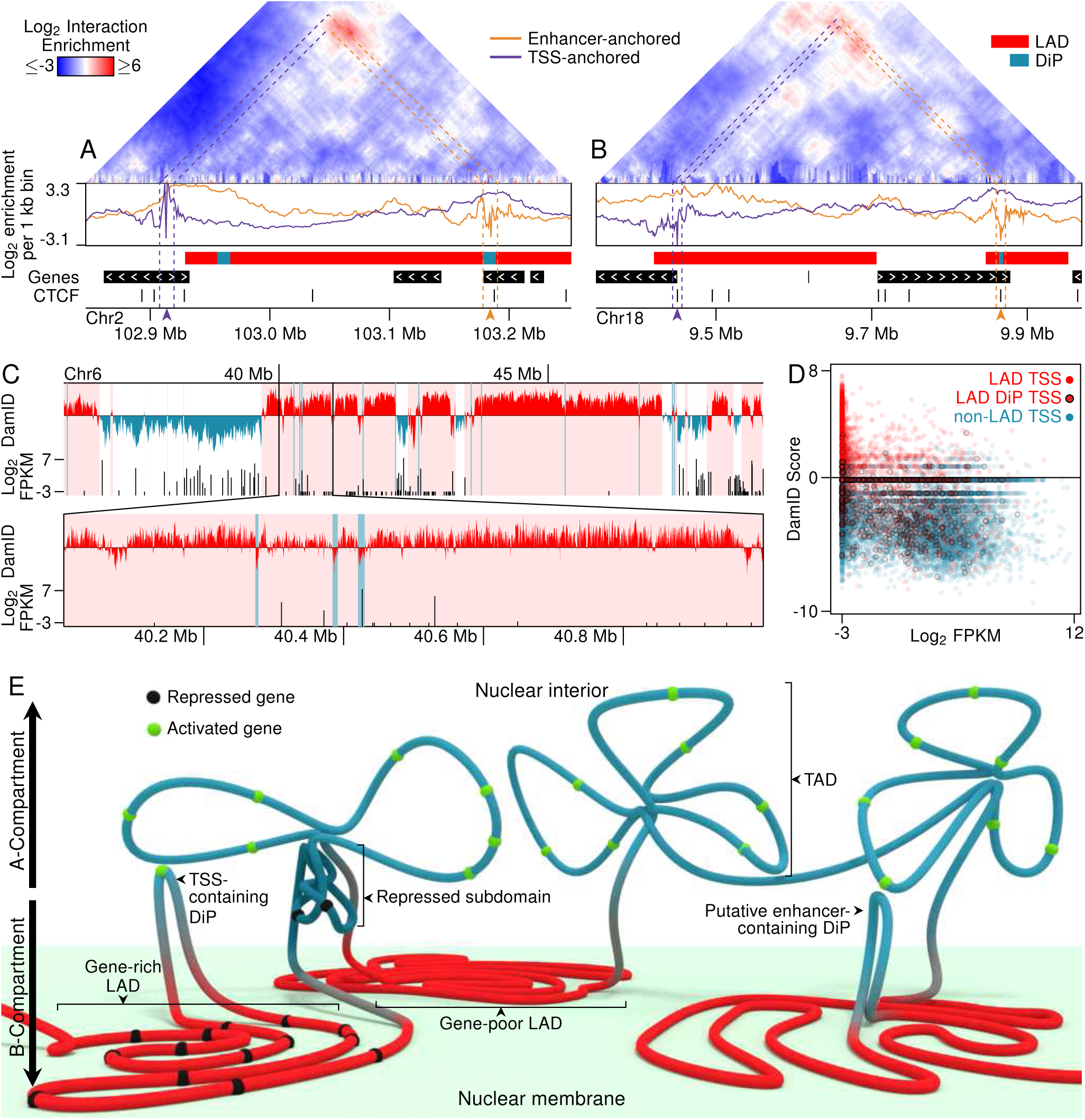
Spatial organization of LADs and their regulatory role. (A) Interaction between PS putative enhancer and potential gene target. Dynamically-binned MEF HiC enrichment data (at least 10 reads per 1 kb bin) show a peak near the intersection of a putative enhancer identified in a LAD DiP and the TSS of the *Apip* gene. A 4C-like plot shows interaction intensities anchored at the putative enhancer (purple) and the gene TSS (orange). (B) Similar peak near the intersection of a putative enhancer identified in a DiP and the TSS of the *Cetn1* gene. (C) A sample region for mouse chromosome 6 in MEFs. LAD calls are highlighted in pink and indicated by red DamID scores. log_2_ transformed gene expression levels (FPKM) are given with unexpressed genes plotted at -3. Blue lines denote LAD DiPs. (D) Genes plotted by log-transformed expression and DamID score associated with their TSS. A minimum score of -3 was used for expression values. Points are color-coded by whether the TSS occurs in a LAD (red) or not (blue). TSSs occurring in LAD DiPs are outlined in black. (E) A cartoon schematic of spatial organization of chromatin structures.

Taking these new data together we propose a framework for interrelationship between LADs and overall chromosome structure (Figure 8E). We determined that, contrary to previous findings, LADs and non-LADs segregate consistently in single cells, both by association to the nuclear lamina but also by the aggregation of individual LADs. This segregation is dependent upon chromatin state and an intact nuclear lamina. In addition, using a refined algorithm to detect higher-order chromatin interactions, we determine that LADs represent the B-compartment, previously shown to harbor transcriptionally silent regions of the genome and implied to be enriched in LADs. However, our analyses also show that regions within the A/B-compartments can be smaller than previously defined, allowing much smaller regions of the genome to be differentially compartmentalized. In general, we find that LADs show relatively low interaction within LADs, in contrast with non-LAD regions, which display robust intra-domain interactions. While LADs represent relatively large domains that interact within the peripheral zone at the nuclear lamina, there are small regions which can loop out and interact with the A-compartment. These DiPs are enriched for TSSs and enhancers and at least some of these interact preferentially with other regulatory elements in the genome.

## Discussion

In this study, we describe the use of high-resolution chromosome conformation paints derived from genome-wide maps of lamina association to uncover basic principles of organization of entire chromosomes *in situ*. We integrate both DamID and Hi-C data from MEFs and pro-B cells to demonstrate that the LADs correspond to the B-compartment, and that the dynamic reorganization of lamina associated domains is also reflected in their compartment assignment. Using the chromosome conformation paints, we demonstrate that LAD (and B-compartment) organization is dependent upon both chromatin state and Lamin A/C. Finally, by interrogating and integrating LAD organization data and Hi-C data from the same cells, we demonstrate that small regions within lamina proximal domains interact not with the lamina, but instead with the active A-compartment. These DiPs escape the repressive regime at the peripheral zone and are enriched for TSSs and active enhancers.

Organization of LADs has been studied in a variety of cell types, both by genome-wide data and by cytological measures. Existing genome-wide approaches allow for global measurement of lamin association with high-resolution, but they determine the average state over a large population of cells, and thus do not capture single cell variability. Single-cell DamID, while capturing some potential single cell variability, is necessarily low resolution (Kind et al., 2015). At the other extreme, cytological measures have lower resolution and typically have only examined a small number of loci simultaneously or small sub-domain regions, but these assays do permit actual in *situ* measurements of locus partitioning within the nucleus. To bridge this gap, we designed a new tool, chromosome conformation paints, to examine chromosome-wide domain configuration *in situ* in primary early pass MEFs and *ex vivo* expanded pro-B cells. Chromosome conformation paints enabled us to determine the organization of LADs and non-LADs, relative to each other and the nuclear lamina, in the context of the entire chromosome in *situ* (Figure 1). We find that LADs are not strictly stochastic in their association with the nuclear lamina, but instead display a constrained configuration. These data seem to be in contrast with previous findings suggesting that LADs are highly stochastic in their association with the nuclear lamina. In this study, we measure LAD organization in early pass primary cells, as opposed to sub-cloned cancer-derived cell lines. In addition, we measure all LADs and non-LADs within an entire chromosome, thus enabling us to count regions away from the lamina as contiguous and constrained within the peripheral zone, regions which, if measured as single point probes, would likely be scored as away from the lamina. These chromosome conformation paints therefore demarcate a zone in which LADs are constrained. Use of these chromosome conformation paints also show that LADs display a large degree of self-association (LAD-to-LAD interactions) over large distances within a chromosome and that LADs display a compact organization compared to non-LAD regions (Figure 1). Both constraint at the nuclear lamina and self-association of LADs depend upon chromatin state (H3K27me3 and H3K9me2/3) and Lamin A/C (Figure 3 and 4).

Given the organization observed in single cells, we sought to understand how this organization related to chromosome structure interrogated using chromosome conformation capture (Hi-C). We developed an improved compartment calling algorithm to demarcate A/B-compartment transitions at higher resolution than previously reported. Using this approach, we determined that LADs are the B-compartment (Figure 2). The boundary regions delimiting these domains, both at long-range (A/B-compartment) and by local neighborhood measures (boundary score), show consistent alignment between Hi-C data and LADs. The boundaries are enriched in known boundary factors, such as CTCF and cohesin. Given that LADs are the B-compartment, then, it was not surprising that the Hi-C data support our in *situ* findings of self-association between LADs (Figure 6). Indeed, by comparing MEFs and pro-B cells and focusing on the *Igh* locus, we uncover a general principal of the interactions of LAD regions. In MEFs, the *Igh* locus is in a LAD and is constrained at the nuclear lamina; it displays low levels of local interactions within the LAD region/compartment, but does interact with more distal LADs (LAD-to-LAD). This is supported by previous findings using 3D-ImmunoFISH wherein the *Igh* locus has been shown to be lamina-proximal in non-B cells and in a locus-extended configuration—with distal and proximal regions of the locus, which is located within a single LAD, displaying low levels of interaction. This is in contrast to what is observed in pro-B cells, where the locus is more centrally located in the nucleus and the distal and proximal segments are more often found in close proximity. While this locus likely displays an extreme example of this type of reconfiguration, we find that our data generally support the idea that chromatin at the lamina is globally more compact, but at a more local scale (within an individual LAD) the chromatin is in an extended configuration, disallowing local looping interactions (Figure 8E). In the Hi-C data, this is reflected as low signal locally and a lack of sub-TAD organization with LADs. These data are also in agreement with a recent report by Boettiger et al. (2016), in which repressed chromatin enriched in polycomb group displays different compaction ratios at different length scales.

Even though LADs generally lack a strong TAD structure/substructure, we find that regions of LADs appear to escape interactions with the nuclear lamina and are enriched for TSSs, supporting a similar finding in H3K9me2/3 domains in which similar regions, termed “euchromatic islands”, were noted (Wen et al. 2012). We term these regions DiPs (Disrupted in Peripheral signal) as they appear to dip away from the nuclear periphery (Figure 8C). In general, the DiPs are enriched in H3K27ac and occupancy by CBP, suggesting that these regions represent active enhancers and/or TSSs. The majority of these TSSs are transcriptionally active (GRO-seq) with relatively low abundance of stable transcripts (RNA-seq). However, the majority of the DiPs do not contain annotated TSSs (Figure 7), but instead display characteristics of active enhancers, including a high ratio of H3K4me1/H3K4me3. Intriguingly, many of the TSSs and putative enhancers are bound by cohesin and CTCF, indicating that these regions may represent stabilized structures “looping out”from the nuclear lamina. To determine if these structures remain in the B compartment, we employed a local compartment analysis of the Hi-C data. Indeed, the DiPs are more A-compartment proximal, supporting our hypothesis that these regions actively loop away from the lamina, even showing discrete enhancer-promoter interactions across LAD boundaries (Figure 8). These data highlight the power of integrating disparate data types, as this fine structure of A/B-compartment looping was not immediately evident from Hi-C analyses alone, but required DamID data to highlight specific regions of interest. By focusing on these regions and performing local compartment analysis it is evident that the A/B-compartment switching can occur over much smaller genomic distances than previously thought.

Our data demonstrate that segregation of the genome into LAD/non-LAD or A/B-compartments is recapitulated *in situ* at the single cell level. These data are consistent with theoretical projections from models based on single-cell Hi-C (Stevens et al., 2017). Current models of genome organization using Hi-C data have focused on two modes of organization: the topologically associating domains (TADs) and A/B-compartments. Generally, these modes of organization have been depicted as hierarchical, that is several TADs assemble to form an A/B-compartment region. This suggests that the A/B-compartment is dependent upon TAD structures. However, more recent findings have demonstrated that A/B-compartments can form in the absence of TADs, indicating that TAD and A/B-compartment organization represent independent layers of organization (Nora et al., 2017; Schwarzer et al., 2016). Intriguingly, in the absence of cohesin, which is required for proper TAD organization, A/B-compartments appear more refined in Hi-C data and correlate with chromatin state. Our data help to refine these models by incorporating the relationship between genome “compartmentalization” as determined by Hi-C data with “nuclear compartmentalization” of the genome as determined by association with nuclear structures, namely the nuclear lamina. Our data demonstrate that LADs are the B-compartment and are constrained at the nuclear lamina in *situ*. This constraint appears to require H3K9me2/3 and H3K27me3 as well as Lamin A/C. Loss of either Lamin A/C or heterochromatin leads to some loss of lamina association and, even more strikingly, LAD-to-LAD interactions, suggesting a loss of the A/B-compartments. These data suggest that LAD organization is an important driving force for functional compartmentalization of the genome.

## Author Contributions

T.R.L., K.L.R. designed the project, T.R.L., X.W., M-C.G., K.L.R. executed the experiments, P.T., R.A.A., K.P., and N.A.Y. designed the oligonucleotide pools for paints, M.E.G.S., T.R.L., X.W. and J.T. analyzed the data, K.L.R. wrote the manuscript with input from J.T., M.E.G.S., and T.R.L. All authors edited and approved the final manuscript.

## Acknowledgements

Thanks to the Reeves lab for help making MEFs; Rakel Tryggvadottir, Colin Callahan, and Sinan Ramazanoglu for help with sequencing; Victoria Hoskins, Michelle Halstead, Jevon Cutler and the members of the Reddy and Taylor labs for useful discussion and feedback. This work was funded in part through the JHU Catalyst and Discovery Awards; T.R.L. was funded from NIH Training Grant T32 GM007445; K.L.R and M-C.G. were funded in part by NIH/NIA grant R21 AG050132; and M.E.G.S and J.T. were funded in part by NIH/NIDDK grant R24 DK106766 and NIH/NHGRI grant U41 HG006620.

## Methods

### Contact for reagent and resource sharing

Further information and requests for resources and reagents should be directed to and will be fulfilled by the Lead Contact, Karen Reddy.

### Experimental models

#### Generation and maintenance of primary murine embryonic fibroblast (MEFs)

For primary MEFs, wild-type eight-week-old C57BL/6 mice were bred and embryos were harvested at E13.5. Individual embryos were homogenized using a razor blade, and cells were dissociated in 3 mL 0.05% trypsin for 20 min at 37°C. Next 2 mL of 0.25% trypsin was added and cells were incubated again at 37°C for 5 min. Cells were pipetted vigorously to establish single cells, passed through a 70 μm cell strainer, pelleted and then plated in 10 cm dishes and labeled as P0. MEFs were cultured DMEM High Glucose with 10% FBS, penicillin/streptomycin, L-glutamine and non-essential amino acids. Cells were cultured for no longer than 5 passages before harvesting for experiments. For initial DamID experiments, longer term-culture C57BL/6 MEFs were purchased from ATCC (American Tissue Culture Collection, CRL-2752) and cultured according to their established protocols, in medium containing DMEM High, 10% FBS, Penicillin/Streptomycin and L-glutamine.

#### Murine pro-B cell culture and maintenance

*Ex vivo* expanded Rag2^-/-^ pro-B cells (Harr et al., 2015) were co-cultured with Op-9 stromal cells (ATCC CRL-2749) in Opti-MEM supplemented with 5% FBS, penicillin/streptomycin, L-glutamine, 0.1% beta-mercaptoethanol and 3 ng/mL interleukin-7 (IL-7), as previously described (Medina et al., 2004; Nakano, 1996).

### Method Details

#### Drug treatments

Primary MEFs were cultured as described and were treated with epigenetic modifying drugs for 24-60 hours, as previously described. Drugs were added to the media at the following concentrations and refreshed at 24 hour intervals: 40 ng/mL TSA (Sigma, 1952), 0.5 μM BIX01294 (Ryan Scientific, RYS-AF-0051), 0.25 μM DZNep (Cayman Chemical, 13828, batch 0443536-5). For 3D-immunoFISH experiments, MEFs were treated with inhibitors while grown on slides. For drug treatment combined with DamID, primary MEFs were treated for 18-24 hours with the specified inhibitor, prior to infection with DamID virus.

#### Lamin A/C knockdown

shRNA-mediated LmnA/C knockdown was carried out as described previously (Harr and Reddy, 2016). Briefly, virus for knockdowns was generated in HEK 293T/17 cells (ATCC CRL-11268) by co-transfecting VSV-G, delta 8.9, and shLmnA/C (Sigma, clone NM_001002011.2-901s21c, 5′-GCGGCTTGTGGAGATCGATAA-3′) or shluciferase (5′-CGCTGAGTACTTCGAAATGTC-3′) with Fugene 6 transfection reagent (Promega E2691). 10 mM sodium butyrate was then added to the transfected cells 3 hours post transfection for an overnight incubation at 37°C, 5% CO_2_. The transfection media containing sodium butyrate was removed the following day and the cells were washed with 1X PBS. Opti-MEM was then added back to the cells which were then incubated at 37°C, 5% CO_2_. Viral supernatant was collected every 12 hours up to 3 collections and the supernatant of all 3 collections were pooled. Primary MEFs were cultured as described and incubated overnight with shLmnA/C or shluciferase fresh viral supernatants supplemented with 4 μg/mL polybrene and 10% FBS for 12-14 hours. Fresh MEF media was then added to the cells after the virus was removed and selected with 10 μg/ml blasticidin. For DamID profiling, cells were infected with DamID virus 4 days post shRNA transduction and cultured for additional 48 hours.

#### DamID Protocol

DamID was performed as described previously (Harr and Reddy, 2016; Vogel et al., 2007; Zullo et al., 2012). Cells were either transduced with murine retroviruses or with lentiviruses harboring the Dam constructs. Self-inactivating retroviral constructs pSMGV Dam-V5 (Dam-Only) and pSMGV Dam-V5-LaminB1 (Dam-LaminB1) were transfected using Fugene 6 transfection reagent (Promega, E2691) into the Platinum-E packaging line (Cell Biolabs, RV-101) to generate infectious particles. These viral supernatants in DMEM were used to directly infect MEF lines. Lentiviral vectors pLGW-Dam and pLGW Dam-LmnB1 were co-transfected with VSV-G and delta 8.9 into HEK 293T/17 packaging cells using the Fugene 6 transfection reagent in DMEM High glucose complete media (DMEM High glucose supplemented with 10% FBS, Penicillin/Streptomycin, L-glutamine). 10 mM sodium butyrate was added to the transfected cells 3 hours post-transfection and left overnight. The following day this media was removed and the cells were washed briefly with 1X PBS before Opti-MEM media was added. Supernatants containing viral particles were collected every 12 hours between 36-72 hours after transfection, and these collections were pooled, filtered through 0.45 μM SFCA or PES, and then concentrated by ultracentrifugation. For infection with retrovirus or lentivirus, MEFs were incubated overnight with either Dam-only or Dam-LmnB1 viral supernatant and 4 μg polybrene. Pro-B cells were “spinfected” for 2 hours at 2000 × g with viral supernatants and 8-10 μg/mL polybrene, recovered for 2 hours, as previously described. Cells were allowed to expand for 2-4 days then pelleted for harvest.

DamID infected MEFs were collected by trypsinization and DNA was isolated using QIAamp DNA Mini kit (Qiagen, 51304), followed by ethanol precipitation and resuspension to 1 μg/ul in 10 mM Tris, pH 8.0. Digestion was performed overnight using 0.5-2.5 μg of this genomic DNA and restriction enzyme DpnI (NEB, R0176) and then heat-killed for 20 minutes at 80°C. Samples were cooled, then double stranded adapters of annealed oligonucleotides (IDT) AdRt (5-CTAATACGACTCACTATAGGGCAGCGTGGTCGCGGCCGAGGA-3) and AdRb (5-TCCTCGGCCG-3) were ligated to the DpnI digested fragments in an overnight reaction at 16°C using T4 DNA ligase (Roche, 799009). After incubation the ligase was heat-inactivated at 65°C for 10 minutes, samples were cooled and then digested with DpnII for one hour at 37°C (NEB, R0543). These ligated pools were then amplified using AdR_PCR oligonucleotides as primer (5-GGTCGCGGCCGAGGATC-3) (IDT) and Advantage cDNA polymerase mix (Clontech, 639105). Amplicons were electrophoresed in 1% agarose gel to check for amplification and the size distribution of the library and then column purified (Qiagen, 28104). Once purified, material was checked for LAD enrichment via qPCR (Applied Biosystems, 4368577 and StepOne Plus machine) using controls specific to an internal Immunoglobulin heavy chain (*Igh*) LAD region (J558 1, 5-AGTGCAGGGCTCACAGAAAA-3, and J558 12, 5-CAGCTCCATCCCATGGTTAGA-3) for validation prior to sequencing (Harr et al., 2015).

#### DamID-seq Library Preparation and Sequencing

In order to ensure sequencing of all DamID fragments, post-DamID amplified material was randomized by performing an end repair reaction, followed by ligation and sonication. Briefly, 0.5-5 μg of column purified DamID material (from above) was end-repaired using the NEBNext End Repair Module (NEB E6050S) following manufacturer’s recommendations. After purification using the QIAquick PCR Purification Kit (Qiagen, 28104), 1μg of this material was then ligated in a volume of 20 μL with 1μl of T4 DNA ligase (Roche, 10799009001) at 16°C to generate a randomized library of large fragments. These large fragments were sonicated (in a volume of 200μL, 10mM Tris, pH 8.0) to generate fragments suitable for sequencing using a Bioruptor^®^ UCD-200 at high power, 30 seconds ON, 30 seconds OFF for 1 hour in a 1.5 mL DNA LoBind microfuge tube (Eppendorf, 022431005). The DNA was then transferred to 1.5 ml TPX tubes (Diagenode, C30010010-1000) and sonicated for 4 rounds of 10 minutes (high power, 30 seconds ON and 30 seconds OFF). The DNA was transferred to new TPX tubes after each round to prevent etching of the TPX plastic. The sonication procedure yielded DNA sizes ranging from 100-200 bp. After sonication, the DNA was precipitated by adding 20 μl of 3M sodium acetate pH 5.5, 500 μl ethanol and supplemented with 3 μl of glycogen (molecular biology grade, 20 mg/ml) and kept at -80°C for at least 2 hours. The DNA mix was centrifuged at full speed for 10 min to pellet the sheared DNA with the carrier glycogen. The pellet was washed with 70% ethanol and then centrifuged again at full speed. The DNA pellet was then left to air dry. 20 μl of 10 mM Tris-HCl was used to resuspend the DNA pellet. 1 μl was quantified using the Quant-iT PicoGreen dsDNA kit (Invitrogen, P7589). Sequencing library preparation was performed using the NEBNext Ultra DNA library prep kit for Illumina (NEB, E7370S), following manufacturer’s instructions. Library quality and size was determined using a Bioanalyzer 2100 with DNA High Sensitivity reagents (Agilent, 5067-4626). Libraries were then quantified using the Kappa quantification Complete kit for Illumina (KK4824) on an Applied Biosystems 7500 Real Time qPCR system. Samples were normalized and pooled for multiplex sequencing.

Reads were sequenced with an Illumina HiSeq2500 sequencer, v4 chemistry, using single-end 100 bp read length according to the manufacturer’s protocol with 1% PhiX spike-in.

#### LAD and non-LAD chromosome-wide probe design and labeling

LADs from murine embryonic fibroblasts were defined through the LADetector algorithm v1 on previously published data using default parameters (Harr et al., 2015;Table S1). Non-LADs were defined as complementary regions to LADs. Centromeres were excluded, and LAD and non-LADs were repeat masked. Probes were selected for chromosomes 11 and 12 *in silico* based on TM and GC content, and those with high homology to off target loci were specifically removed. 150 base pair oligos were chemically synthesized using proprietary Agilent technology and probes were labeled in either Cy3 or Cy5 dyes using the Genomic DNA ULS Labeling Kit (Agilent, 5190-0419). 40 ng of LAD and non-LAD probes were combined with hybridization solution (10% dextran sulfate, 50% formamide, and 2X SSC) then denatured at 98°C for 5 minutes and pre-annealed at 37°C.

#### *Igh* BAC probe generation

The CT7-562A21 (Invitrogen) BAC clone, specific for a distal region in the *Igh* locus, was labeled with Dig during nick translation using the DIG-NICK Translation Mix, (Roche, 11745816910) followed by purification using QIAquick PCR Purification spin columns (Qiagen, 28104; (Kosak et al., 2002). This probe was resuspended in a hybridization solution (2X SSC, 10% dextran sulfate,50% Formamide) with blocking DNAs (9 μg salmon sperm DNA,6 μg placental DNA, 3 ug COT-1 DNA) and hybridized according to the 3D-immunoFISH protocol.

#### 3D-ImmunoFISH and immunofluorescence

3D-immunoFISH was performed as described previously (Harr et al., 2015; Irina Solovei, 2010; Reddy and Singh, 2008; Reddy et al., 2008). Briefly, primary fibroblast cells were plated on poly-L-lysine coated slides overnight. pro-B cells were prepared similarly, but plated on slides for 15-30 minutes in a humid chamber at 37°C and mildly hypotonically treated in 0.3X PBS for 30 seconds to prevent shrinking in the paraformaldehyde fixation. Cells on slides were fixed in 4% paraformaldehyde (PFA)/1X PBS for 12 minutes (pro-B cells) or 15 minutes (fibroblasts and MEFs), then subjected to 3-5 minute washes in 1X PBS. After fixation and washing, cells were permeabilized in 0.5% TritonX-100/0.5% saponin for 15-20 minutes. The cells were washed 3 times 5 minutes each wash in 1X PBS, then acid treated in 0.1N hydrochloric acid for 12 minutes at room temperature. After acid treatment, slides were placed directly in 20% glycerol/1X PBS and then incubated at least one hour at room temperature or overnight at 4°C. After soaking in glycerol, cells were subjected to 4 freeze/thaw cycles by immersing glycerol coated slides in a liquid nitrogen bath. Cells were treated with RNAse (100 μg/ml) for 15 min in 2X SSC at room temperature in a humidified chamber. DNA in cells was denatured by incubating the slides in 70% formamide/2X SSC at 74°C for 3 min, then 50% formamide/2X SSC at 74°C for 1 min. After this denaturation, cells were covered with a coverslip containing chromosome conformation paints or BAC-derived probes in hybridization solution and sealed. After overnight incubation at 37°C, slides were washed three times in 50% formamide/2X SSC at 47°C, three times with 63°C 0.2X SSC, one time with 2X SSC, and then two times with 1X PBS before blocking with 4% BSA in PBS for 30-60 min in a humidified chamber. Slides were then incubated with primary antibody (1:200 dilution; Santa Cruz, SC-6217) in blocking medium overnight at 4°C. Slides were washed three times with 1X PBS/0.05% Triton X-100 and then incubated with secondary antibody in blocking medium DyLight 488 (1:200 dilution; Jackson ImmunoResearch, 211-482-171) for 1 hour at room temperature. Post incubation, slides were washed three times with 1X PBS/0.05% Triton X-100, and then DNA counterstained with 1 μg/ml Hoechst. Slides were then washed, mounted with SlowFade Gold (Life Technologies, S36936).

### Quantification and statistical analysis

#### Image acquisition and processing

Slides were imaged using a Zeiss Axiovert fitted with an ApoTome and AxioCam MRm Camera. Imaging was performed at 100x or 63x with an Apochromat oil immersion objective with an NA of 1.5 using Immersol 518. AxioVision software was used to acquire images and .zvi files were exported and processed in FIJI (FIJI is just ImageJ; (Schindelin et al., 2012). Chromosome territories were evaluated for nuclear position and attachments to the lamina. To quantify the degree of association with the nuclear lamina, chromosomal subdomains were measured by taking the average of three lines through each territory in the medial planes of the image and plotting the distance from the peak Lamin B1 signal for LAD and non-LAD domains (Yao et al., 2011). The distribution of LAD and non-LAD signals was measured using line scans in triplicate from outside to inside the nucleus and histogram measurements of pixel intensity were acquired for each channel using FIJI. Nuclei that were polyploid for chromosome 11 or 12, exhibited damage or were not fully visible in the field were excluded from the analysis. For each measurement, maximum lamin B1 signal was set to x=zero and all distances are relative to this zero point. Distance measurements from a single chromosome (three independent measurements) were averaged together and normalized by total pixel intensity (Normalized value = Pixel intensity/Sum of total pixel intensity). Data was collected from multiple experiments performed on different days and results were pooled.

#### DamID-seq Data Processing

DamID-seq reads were processed using LADetector (https://github.com/thereddylab/LADetector), an updated and packaged version of the circular binary segmentation strategy previously described for identifying LADs from either array or sequencing data (Harr et al., 2015). LADetector relies on several programs for analysis steps: Bowtie v2.2.3 (Langmead &Salzberg,2012), R v3.2.0 (R Core Team, 2015), BedTools v2.19.1 (Quinlan & Hall, 2010), and SamTools v1.3 (Li et al., 2009). For arrays, DamID array signal intensity data quantile normalized, smoothed with the preprocessCore R package (Bolstad, 2003), and analyzed using LADetector v1 and merged. Sequencing data were analyzed using LADetector v2 and LADs and DiPs were output. DiPs (Disruption in Peripheral signal) size was set to 1-25 kb; LADs separated by less than 25 kb were considered to be part of a single LAD. LADs were post-filtered to be greater than 100 kb, complementary genomic regions to LADs were defined as non-LADs. BedGraphs were generated for visualization using bedtools genomecov. Scripts, files and workflow for DamID analysis are available at https://github.com/thereddylab/LADetector.

#### ChIP-Seq Data Processing

ChIP-seq sequencing data and associated controls were downloaded from SRA (Supplemental Table 1). Data were aligned to the mouse genome build 9 using Bowtie version v1.1.1 (Langmead et al., 2009) allowing up to 2 seed mismatches, the `“–tryhard”` option, and only reporting uniquely mappable reads. All other parameters were left with default values. Peak calls and pileup tracks were generated using MACS2 version 2.1.1.20160309 (Zhang et al., 2008). With the exception of setting the genome to mouse, all other parameters were left with default values. For histone marks, the `“–broadflag”` was used.

#### Hi-C normalization

Raw sequences for MEF and pro-B cells (Supplemental Table 1) were obtained from SRA (Krijger et al., 2016; Lin et al., 2012). Read ends were aligned to the mouse genome build mm9 using BWA mem version 0.7.12-r1039 and default settings (Li and Durbin, 2009). Reads were kept if they met one of the following criteria: Each read end mapped to a single position; one end failed to map but the other end mapped to two positions falling in two different restriction fragments; both ends mapped to no more than two positions from different restriction fragments and the downstream position of one end occurred on the same fragment as the upstream position of the other end. For MEF data, all replicates were combined. Reads were processed and normalized using HiFive version 1.3.2 (Sauria et al., 2015). A maximum insert size of 650 bp was used to filter reads. Fends (fragment ends) were filtered to have a minimum of one valid interaction greater than 500 kb.

The data were normalized using the “binning” algorithm correcting for GC content, fragment length, and mappability. GC content was calculated from the 200 bp upstream of restriction sites or the length of the fragment, whichever was shorter. Mappability was determined using the GEM mappability function, version 1.315 (Koehler et al., 2011). Mappability of 36-mers was calculated every 10 bp with an approximation threshold of six, a maximum mismatch of 1 bp, and a minimum match of 28 bp. For each fend, the mean mappability score for the 200 bp upstream of the restriction site, or total fragment size if smaller, was used. For normalization, only intra-chromosomal reads with an interaction distance of at least 500 kb were used. GC content and fragment length were partitioned into 20 bins each and mappability was partitioned into 10 bins. All parameter partitions were done such that together they spanned the full range of values and contained equal numbers of fends in each bin. All bins were seeded from raw count means and GC and length parameters were optimized for up to 100 iterations or until the change in log-probability was less than one, whichever was achieved first.

#### Hi-C Compartment Scoring

Eigenvector-based compartment scores were calculated as previously described (Jung et al., 2017). Enrichments were calculated for either 1 Mb (low-resolution) or 10 kb bins (high-resolution). Bins were expanded using HiFive’s dynamic binning to a minimum or 3 reads per bin. For each pairwise combination of rows for the enrichment heatmap, the Pearson correlation was calculated. Taking the first eigenvector of the correlation matrix yielded the eigenvector-based compartment score. Because the sign of the eigenvector is random, we used mean transcriptional activity in positively versus negatively scored regions to determine A and B-compartment score signs. Where necessary, signs were flipped so that all B-compartments corresponded to positive eigenvector scores.

Likelihood compartment scores were calculated as the log_2_-transformed ratio of the probability of each 10 kb interval occurring in the B-compartment divided by the probability of that interval occurring in the A-compartment. The sign of the high-resolution first eigenvector score described above determined compartment initialization (positive values were associated with the B-compartment). Bins with fewer than five interactions longer than 1.5 Mb were removed. Interactions spanning 1.5 Mb or greater were divided into three groups: both sides occurring in the A-compartment, the B-compartment, or one side in each compartment. The distance dependent signal curve for each category was calculated by finding the sum of counts divided by the sum of expected values at each distance interval. For distance, intervals containing fewer than 10000 reads were joined with the next largest interval prior to finding enrichment. This was continued until the 10000 read minimum was met. The effective distance for joined bins was calculated as the mean of the log-transformed bin distances. Enrichment values for distances corresponding to bins that had fewer than 10000 reads were interpolated linearly based on the log-transformed expected values and log-transformed distances of the two adjacent bins. The probability for each interval was calculated under the Poisson distribution as follows:

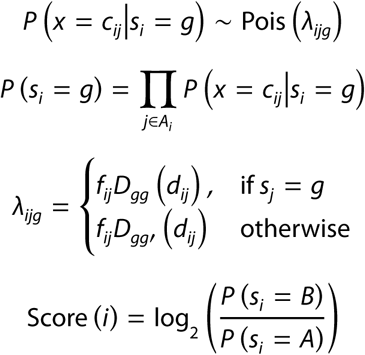
where *s*_*i*_ is the compartment state of interval *i, A*_*i*_ is the set of valid interactions of at least 1.5 Mb involving interval *i, c*_*ij*_ is the sum of observed counts for the interaction bin between intervals *i* and *j, f*_*ij*_ is the sum of interaction normalization values for bin *ij, d*_*ij*_ is the distance between midpoints of intervals *i* and *j*, and *D*_*gg*_ and *D*_*gg′*_ are the distance dependent signal functions for within compartments of type b and between different compartments, respectively.

Training was performed on a chromosome by chromosome basis in an iterative fashion, calculating the distance dependent signal curves, calculating the compartment scores, updating the top 50% of scores (rounding up), and adjusting states based on the signs of the scores. This was performed for up to 200 rounds. If a stable set of interactions was achieved, the associated scores were kept. If a chromosome began switching between two sets of stable states, the mean of these two sets of scores was taken. Otherwise after a 20 round burn-in period, scores were sampled every round and the mean score for each interval was taken after the final iteration.

#### Hi-C Boundary Index Scoring

The Boundary Index score captures shifts in interaction partner preference without the requirement that the majority of interactions be primarily upstream or downstream on either side of boundary points. This is accomplished by finding correlations for 2.5 kb binned reads across 25 kb by 25 kb pools of reads, extending up- and downstream 250 kb and stepping at 2.5 kb intervals, excluding the 50 kb closest to the boundary point. For the genomic position at the start of bin *i*, the boundary scores is calculated as follows:

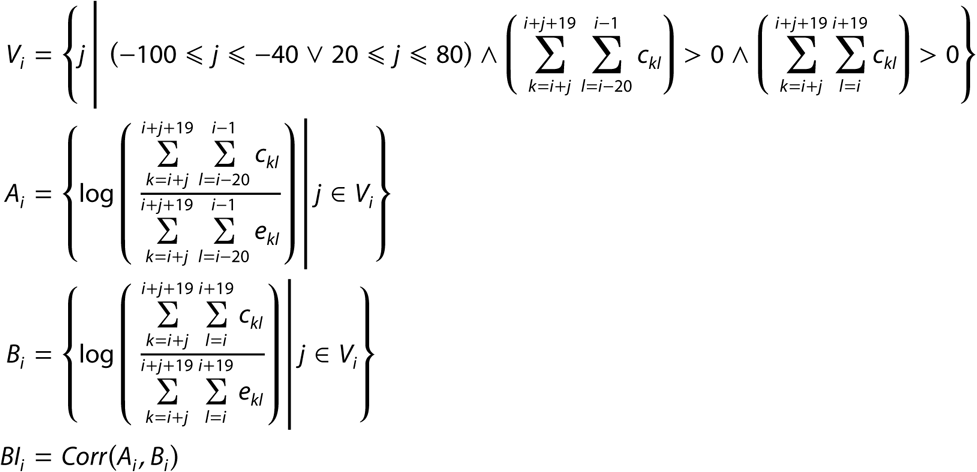
where *c*_*kl*_ and *e*_*kl*_ are the read counts and sum of enrichment expected values for the interaction bin between intervals *k* and *l* under HiFive’s normalization model.

#### Hi-C Chromosome Modeling

For each chromosome, fend-normalized reads were dynamically binned using HiFive with a bin size of 10 kb for both raw and expansion binning and were expanded to ensure a minimum of 3 reads per bin. Enrichments were calculated as the log-transformed ratio of counts to expected values. The diagonal of the enrichment matrix (self-interacting bins) were set to the maximum off-diagonal enrichment value of the matrix. The first three eigenvectors were calculated using the PCAfast function from the python library “mlpy” (Albanese et al., 2012). These three values were used as coordinates for each bin.

#### Domain Interactions

For each pairwise region between intervals bound by LAD boundaries (both between LADs/non-LADs and between LADs and non-LADs), cumulative count and expected values were calculated. LAD calls prior to 100 kb size filtering were used for this analysis. Regions with fewer than 20 reads were discarded.

#### TSS enrichment in DiPs

Enrichment of TSSs in DiPs was calculated by random reordering of the inter-TSS spacing within DiP-containing LADs, maintaining TSS density and number. This was repeated for 100,000 iterations, calculating the number of TSSs falling in DiPs at each iteration.

### Data and software availability

The data discussed in this publication have been deposited in NCBI’s Gene Expression Omnibus (Edgar et al., 2002) and are accessible through GEO Series accession number GSE97095 (https://www.ncbi.nlm.nih.gov/geo/query/acc.cgi?acc=GSE97095).

A “track hub” for visualizing the datasets described in this study in the UCSC genome browser is available at http://genome.ucsc.edu/cgi-bin/hgTracks?db=mm9&position=chr12&hubUrl=https://bx.bio.jhu.edu/track-hubs/icc/hub.txt. The LADetector software is available from https://github.com/thereddylab/LADetector and the HiFive software is available from https://github.com/bxlab/hifive. A companion website with these resources is available at https://bxlab.github.io/conformation-paints-2017/.

